# Sterol-lipids enable large-scale, liquid-liquid phase separation in bilayer membranes of only 2 components

**DOI:** 10.1101/2024.02.02.578692

**Authors:** Kent J. Wilson, Huy Q. Nguyen, Jacquelyn Gervay-Hague, Sarah L. Keller

## Abstract

Despite longstanding excitement and progress toward understanding liquid-liquid phase separation in natural and artificial membranes, fundamental questions have persisted about which molecules are required for this phenomenon. Except in extraordinary circumstances, the smallest number of components that has produced large-scale, liquid-liquid phase separation in bilayers has stubbornly remained at three: a sterol, a phospholipid with ordered chains, and a phospholipid with disordered chains. This requirement of three components is puzzling because only two components are required for liquid-liquid phase separation in lipid monolayers, which resemble half of a bilayer. Inspired by reports that sterols interact closely with lipids with ordered chains, we tested whether phase separation would occur in bilayers in which a sterol and lipid were replaced by a single, joined sterol-lipid. By evaluating a panel of sterol-lipids, some of which are found in bacteria, we discovered a minimal bilayer of only two components (PChemsPC and diPhyPC) that robustly demixes into micron-scale, liquid phases. It suggests a new role for sterol-lipids in nature, and it reveals a membrane in which tie-lines (and, therefore, the lipid composition of each phase) are straightforward to determine and will be consistent across multiple laboratories.

**Significance Statement:** A wide diversity of bilayer membranes, from those with hundreds of lipids (e.g., vacuoles of living yeast cells) to those with very few (e.g., artificial vesicles) phase separate into micron-scale liquid domains. The number of components required for liquid-liquid phase separation has been perplexing: only two should be necessary, but more are required except in extraordinary circumstances. What minimal set of molecular characteristics leads to liquid-liquid phase separation in bilayer membranes? This question inspired us to search for single, joined “sterol-lipid” molecules to replace both a sterol and a phospholipid in membranes undergoing liquid-liquid phase separation. By producing phase-separating membranes with only two components, we mitigate experimental challenges in determining tie-lines and in maintaining constant chemical potentials of lipids.

## Introduction

Lipid bilayers are best known as membranes that surround cells. At low temperatures, lipids are in solid phases. However, in most biological contexts, lipid molecules are in liquid phases: they diffuse freely in the membrane, and their carbon chains are not rigid. When multiple types of lipids are present in membranes, they have the potential to undergo liquid-liquid phase separation. This miscibility phase transition has applications from fundamental biology (e.g., understanding how yeast vacuole membranes acquire domains that allow the cells to survive periods of nutrient reduction (1)) to fundamental physics (e.g., measuring 2-dimensional dynamic critical exponents in systems with conserved order parameter (2)).

Underlying these grand applications is a mystery. Except in extraordinary circumstances (3, 4), all lipid bilayers that have been found to phase separate into large-scale, coexisting liquid phases have been comprised of at least three lipids: a sterol like cholesterol, a lipid with ordered chains, and a lipid with disordered chains. Empirically, this rubric translates into mixtures of a sterol, a lipid with a high melting temperature (*T*_melt_) and a lipid with a low melting temperature (Fig. 1) (5–7).

**Figure 1:**
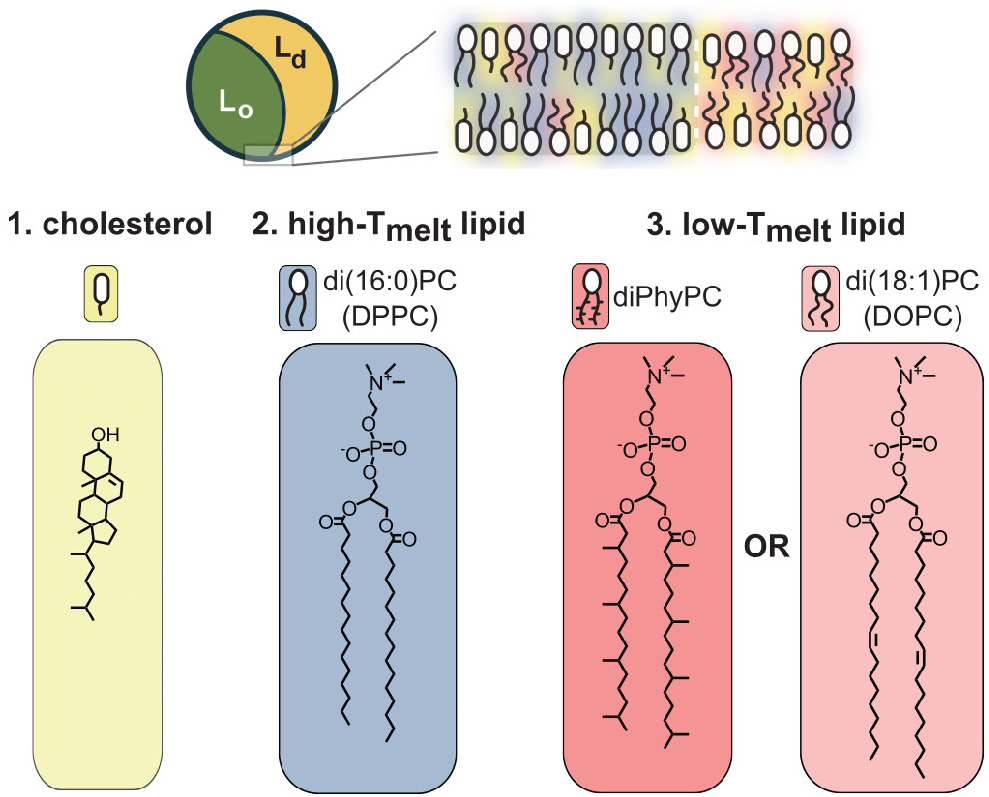
Typically, bilayers that undergo liquid-liquid phase separation contain at least three lipid components. The Lo phase is enriched in high-Tmelt lipid and cholesterol, whereas the Ld phase is enriched in low-Tmelt lipid.

The requirement of *three* components in a lipid bilayer is perplexing. Phase separation of only *two* components is theoretically allowed by the Gibbs Phase Rule in systems where temperature and pressure freely vary. Mean-field theories that consider chain order and interactions give rise to liquid-liquid phase separation in ternary as well as binary membranes (8). Moreover, lipid monolayers (which, at their simplest, are half of a bilayer) require only two components to demix into micron-scale liquid domains (9, 10), even at higher surface pressures (11, 12). Monolayers account for one of the known exceptional circumstances for bilayers: when two monolayers are deposited on a substrate to form a bilayer, domains from each monolayer can persist (3). The other exception arises when cholesterol is chemically altered through the addition of a hydrophobic group on its acyl tail. The resulting hydroxysterol is found in multiple configurations (parallel, anti-parallel, and perpendicular to the other lipids in the bilayer, in crystals, and possibly in hydrogen-bonded clusters) (4). If these configurations interconvert quickly on the time scale of experiments, then all hydroxysterol molecules in the bilayer act as one component. If they interconvert slowly, then each configuration can represent a separate component. Similar ideas hold in systems in which polymorphs of a single type of atom constitute different components (e.g., tetrahedrons vs. polymers of phosphorus, or short-vs. long-chain polymers of sulfur) (13, 14)

There are good reasons to seek a truly minimal, 2-component lipid membrane for liquid-liquid phase separation. Modelers could use it as a test case for simulations. Experimentalists could be confident that membranes produced in their labs demix along the same tie-line and produce phases of the same lipid composition as those in any other laboratory at the same temperature.

If we were to search for binary lipid compositions that phase separate into micron-scale liquid ordered (Lo) and liquid disordered (Ld) phases in bilayers, where would we start looking for them? Lo phases in bilayers are enriched in high-*T*_melt_ lipids and, to a lesser extent, sterols. These molecules experience Van der Waals interactions and have the potential to hydrogen bond if the high-*T*_melt_ lipid is a sphingomyelin (15–17). If membrane phase separation is at least partially driven by close interactions between the sterol and the high-*T*_melt_ lipid, then it is reasonable to surmise that the two molecules could be replaced by a single, joined sterol-lipid, which would decrease the number of components.

To that end, we produce vesicle membranes from binary mixtures of a phospholipid with each of the sterol-lipid molecules in Fig. 2, and we test if the membranes undergo liquid-liquid phase separation to form micron-scale domains. The sterol-lipids represent a range of structures in which a sterol is linked to a hydrocarbon chain. A strong advantage of these structures is that the sterol should be in only one configuration: parallel to the surrounding lipids. The lack of additional configurations of the sterol means that the systems we investigate are truly 2-component.

- In **Sterol-lipid A1 (PChemsPC)**, cholesterol replaces one of the saturated hydrocarbon chains of a phospholipid. A closely related molecule, **Sterol-lipid A2 (PChcPC)**, differs only in the linkage between the sterol and the glycerol backbone.
- **Sterol-lipid B (BbGL-1**) is similar but lacks a phosphate group. Instead, cholesterol and a hydrocarbon chain are β-linked via a galactose moiety. Sterol-lipid B is found in *Borrelia burgdorferi*, the bacterium responsible for Lyme Disease (18).
- **Sterol-lipid C (CPG)**, which is found in *H. pylori*, is composed of a phospholipid α-linked to cholesterol via glucose, a sugar in a ring form (19). Unlike the other sterol-lipids in our set, it has the same 2:1 ratio of acyl chains to sterol group as the two molecules it replaces.
- In **Sterol-lipid D (OChemsPC)**, an unsaturated hydrocarbon chain replaces the saturated chain of Sterol-lipid A. This molecule serves as a control because it has a low-*T*_melt_ carbon chain rather than a high-*T*_melt_ chain.

**Figure 2:**
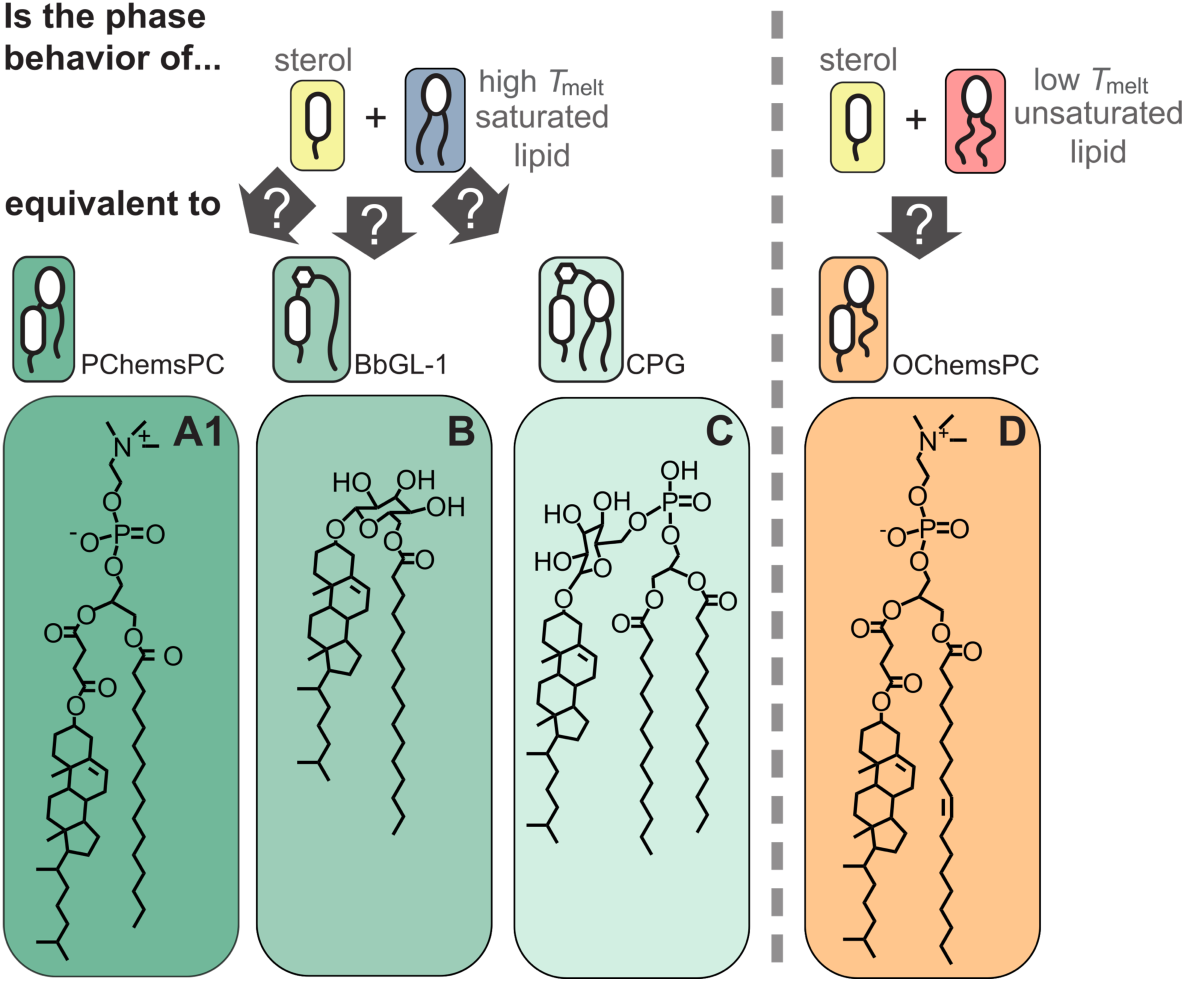
We test if micron-scale, liquid-liquid phase separation occurs in membranes with only two components, one of which is a sterol-lipid. The acyl chains of the sterol-lipids we use are either saturated, as in Sterol-lipid A1, B, and C (known as PChemsPC, BbGL-1, and CPG, respectively) or unsaturated as in Sterol-lipid D (OChemsPC). The acyl chain lengths of Sterol-lipids A1, B, C, and D are 16, 16, 14, and 18, respectively.

It seems easier to speculate about whether a mythical centaur accurately represents both a human and a horse than to predict whether a sterol-lipid can replace both cholesterol and a high-*T*_melt_ lipid in a phase-separating membrane. One reason to be skeptical that binary lipid membranes with sterol-lipids could form micron-scale liquid phases is that most phase-separating membranes contain different ratios of cholesterol to high-*T*_melt_ lipid in their Lo and Ld phases (20), whereas this ratio is necessarily constant in a sterol-lipid. Moreover, it is not clear if membranes containing sterol-lipids should always be stable. From their structures in Fig. 2, sterol-lipids appear to be cone-shaped, with larger tails than headgroups; they may promote curved inverted hexagonal (H_II_) structures rather than flat lamellae. On the other hand, there are also good reasons to hypothesize that binary membranes of phospholipids and sterol-lipids will demix into liquid domains. From a theoretical standpoint, a sterol-lipid could correspond to a complex between cholesterol and a reactive phospholipid (21). From an experimental standpoint, some sterol-lipids have been incorporated into ternary or multi-component membranes at moderate concentrations, where they can substitute for cholesterol and allow micron-scale liquid-liquid phase separation to occur (22). For example, multi-component, phase-separating membranes have been made from mixtures of sterol-lipids (one of which is Sterol-lipid B) and phospholipids or from mixtures of a single sterol-lipid (CAG) with all of the PE lipids extracted from *H. pylori*, which encompass a range of acyl chains (22, 23).

## Results and Discussion

### Robust liquid-liquid phase separation in binary membranes

It is well known that lipid membranes comprised of a sterol, a high-*T*_melt_ lipid, and a low-*T*_melt_ lipid can undergo liquid-liquid phase separation (5–7). Here, we tested whether two of these three components could be replaced by a single, joined sterol-lipid (Fig. 2). Specifically, we hypothesized that we would observe micron-scale, liquid domains in giant vesicles composed of binary mixtures of saturated sterol-lipids and low-*T*_melt_ lipids. All experiments and controls in this study are summarized in Fig. 3 and Fig. 4, respectively.

**Figure 3:**
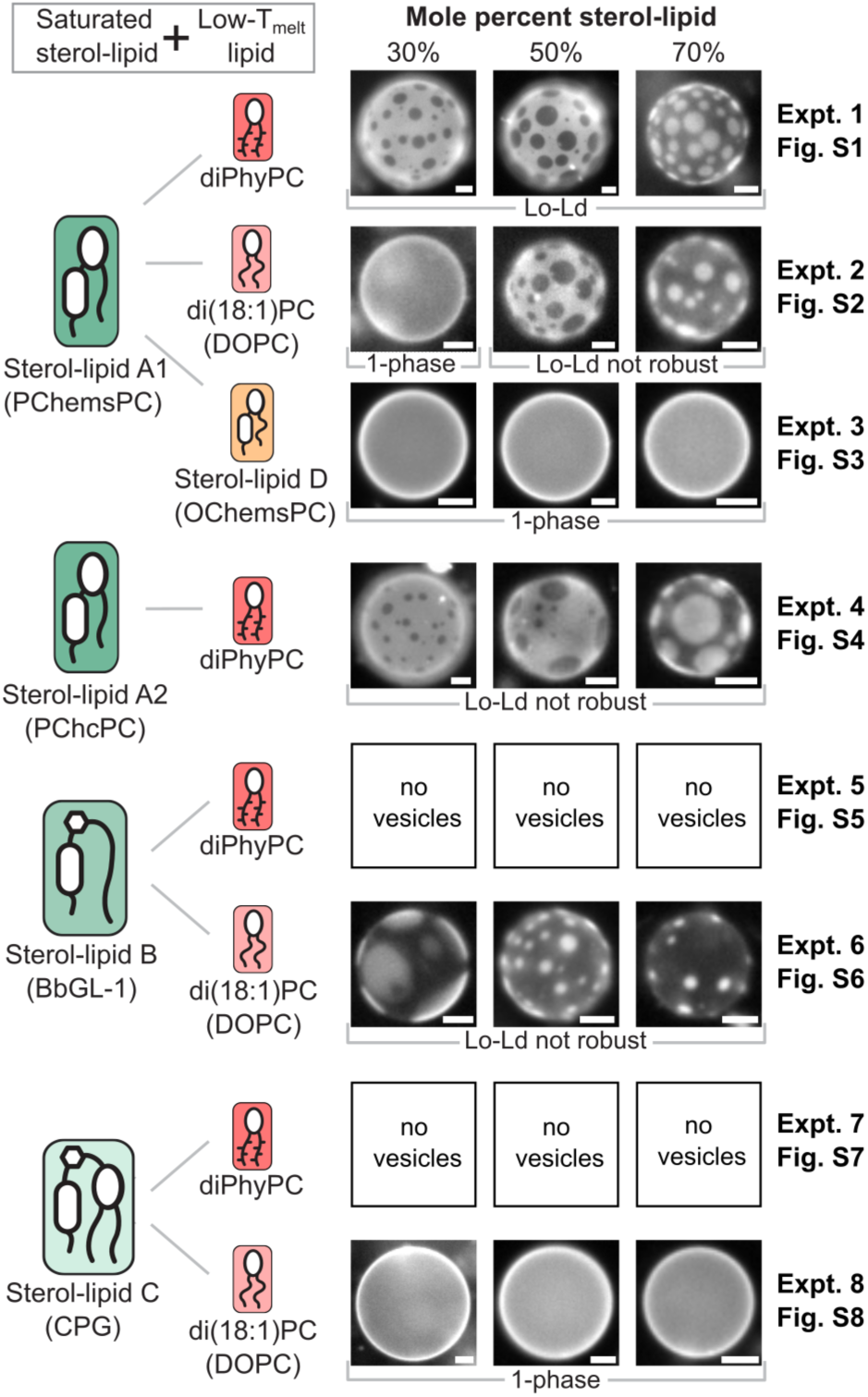
Seven panels of experiments to determine if liquid-liquid phase separation occurs in vesicles composed of binary lipid mixtures in which one component is a saturated sterol-lipid and the other component has a low melting temperature. Experiment 1 is the only case in which liquid-liquid phase separation was observed both with and without a slight osmotic pressure difference across the membrane. The designation “Lo-Ld not robust” means ≤ 50% of vesicles displayed liquid domains only in slightly hyperosmotic solutions, and that no vesicles phase separated without an osmotic pressure difference. Other vesicles displayed only solid-liquid coexistence or one uniform liquid phase. For some combinations, no vesicles formed. All vesicles are shown at 8°C. Scale bars on all fluorescence micrographs are 10 µm. Wide fields of view are in supplementary Fig S1-S8.

**Figure 4:**
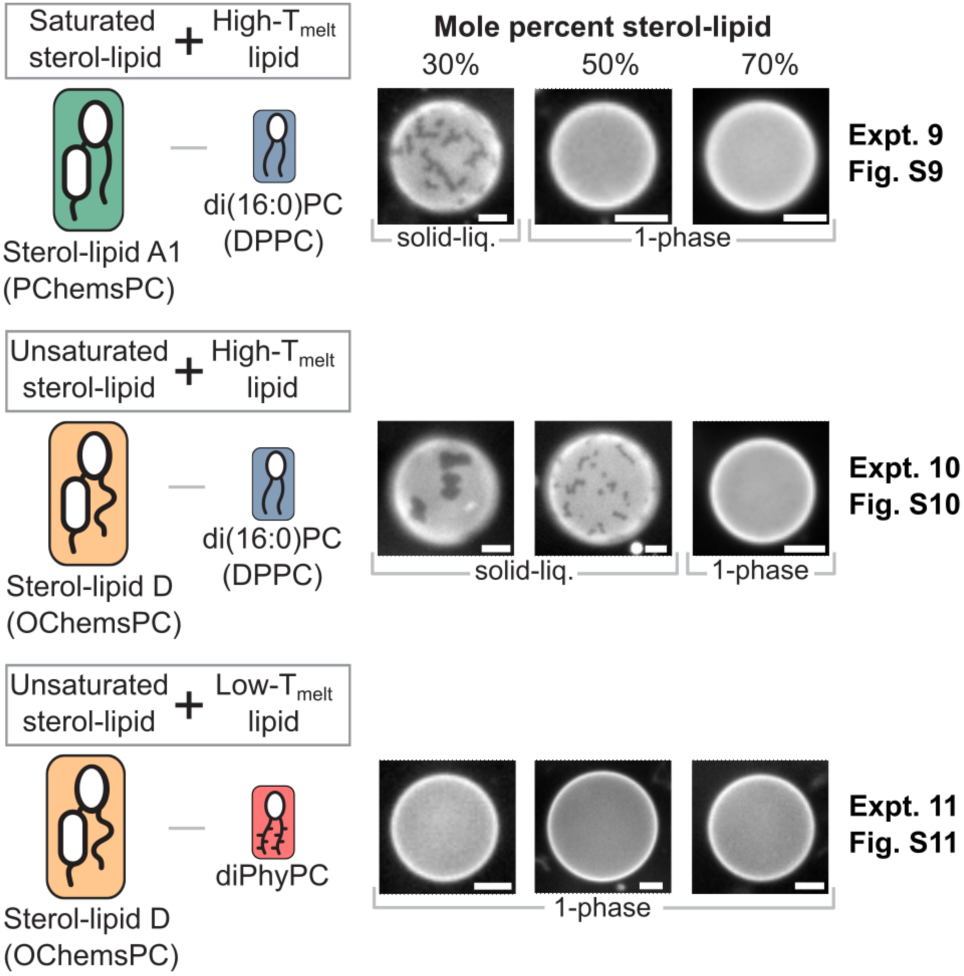
Three panels of negative controls to determine if liquid-liquid phase separation occurs in vesicles composed of binary lipid mixtures that lack a saturated sterol-lipid and/or lack a phospholipid with a low melting temperature. No liquid-liquid phase separation was observed. Instead, vesicles displayed only solid-liquid coexistence or one uniform liquid phase. All vesicles are shown at 8°C. The scale bar on all fluorescence micrographs is 10 µm. Wide fields of view are in supplementary Fig S8-S11.

Our central result, shown in Experiment 1 of Fig. 3, is that binary lipid membranes containing a sterol-lipid do indeed undergo liquid-liquid phase separation. Demixing is most robust when one lipid is saturated Sterol-lipid A1 (PChemsPC) and the other is a phospholipid with a very low *T*_melt_ (diPhyPC). Characteristic of liquid phases, the membrane domains are circular (Fig. 5A), and they coalesce quickly when they collide (Fig. S12 and Movie S2). In taut vesicles, domains merge until only one domain of each phase remains, so domain size is a function of the time elapsed since domains nucleated (24) and tension in the membrane (25). At lower temperatures, nearly 100% of vesicles in Experiment 1 demix. The miscibility transition temperature, *T*_mix_, is defined at the half-max (Fig. 5B).

**Figure 5:**
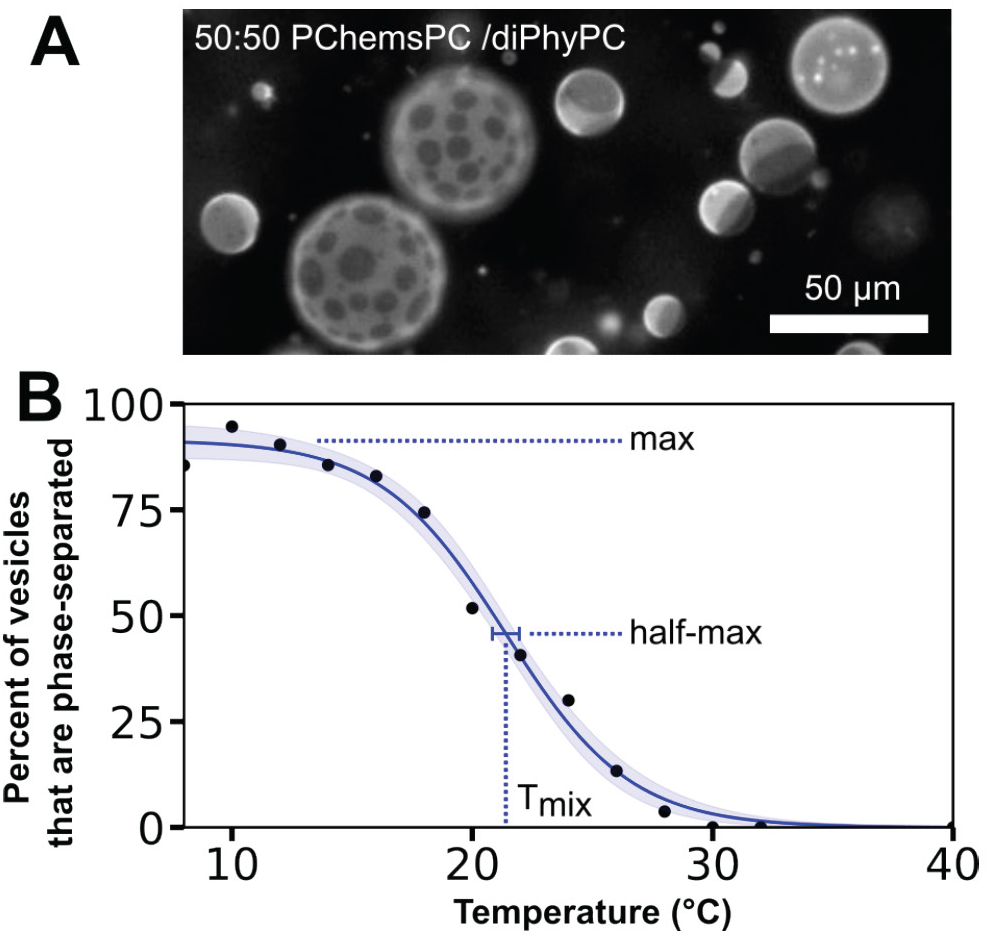
(A) Fluorescence micrograph of micron-scale liquid domains in membranes of vesicles of a 50:50 mixture of a sterol-lipid (PChemsPC) and a low-Tmelt lipid (diPhyPC) at 10°C. Bright regions are Ld phase and dark regions are Lo phase. Vesicles were made in 100 mM sucrose and diluted in 80mM sucrose. (B) Roughly 100 vesicles were observed at each temperature to determine the miscibility transition temperature, Tmix. The shaded region is the 95% confidence interval of the fit.

Our hypothesis was that a single, joined sterol-lipid can behave as a combination of a sterol and a high-*T*_melt_ lipid in enabling phase separation of a bilayer. This hypothesis leads to the expectation that the Lo phase of binary membranes is enriched in sterol-lipid, just as the Lo phase of ternary membranes is enriched in sterols and high-*T*_melt_ lipids (26). Consistent with that expectation, the area fraction of dark, Lo phase increases in vesicles as the mole fraction of sterol-lipid increases (Fig. 3). Avanti Polar Lipids quotes PChemsPC as >99% pure. In many membranes with PChemsPC, small crystallites cover ∼0.1% of the membrane area. (Fig. S13). If the crystallites are due to an impurity, that impurity is too sparse to detect by thin layer chromatography (Fig. S14).

Binary phase diagrams are straightforward to interpret. In Fig. 6, we compile the *T*_mix_ values we measured for binary mixtures of Sterol-lipid A1 and a phospholipid. Several aspects of this phase diagram make it particularly appealing. **1)** The membrane is either in one uniform phase or in two coexisting liquid phases, over the broad range of 20-80 mol% sterol-lipid. **2)** The membrane has a clear upper miscibility critical point. The highest transition temperatures correspond to 30-50 mole% sterol-lipid. Indeed, for membranes near these highest transitions, domain shapes fluctuate, characteristic of low line tension near a critical point (supplementary Fig. S1 and Movie S1) (2). **3)** Binary phase diagrams greatly simplify the task of evaluating tie-lines. Finding the ratio of each lipid in the Ld and Lo phases at 20°C is as easy as following the horizontal grid line across the diagram at 20°C and noting that it intersects the transition to one uniform phase at roughly 30% and 70% sterol-lipid, the compositions of the Ld and Lo phases at that temperature.

**Figure 6:**
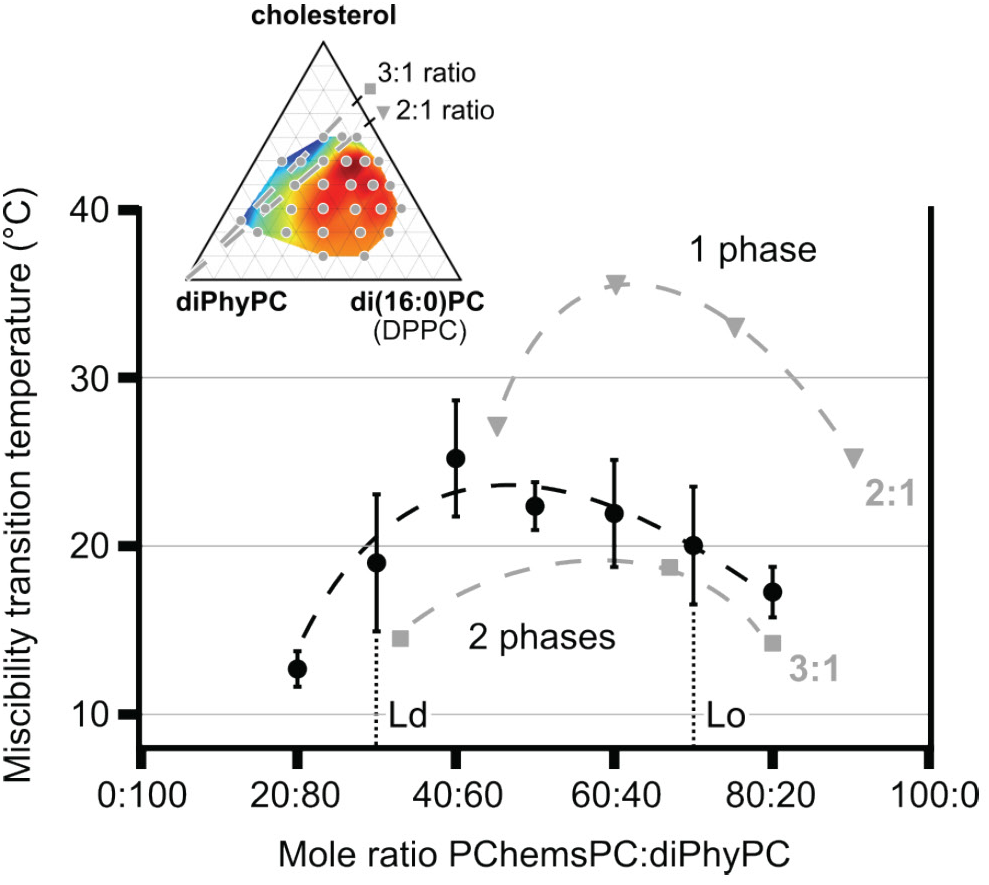
Black circles are miscibility transition temperatures of binary mixtures of Sterol-lipid A1 (PChemsPC) and a low-Tmelt lipid (diPhyPC) in vesicles containing 100 mM sucrose, free-floating in a solution of 80 mM sucrose. Uncertainties are standard deviations from three independent experiments at each lipid ratio. The transition is reversible with temperature. At 20°C, endpoints of the tie-line fall at ratios of roughly 30:70 and 70:30 sterol-lipid to high-Tmelt lipid, the compositions of the Ld and Lo phases. Membranes at ratios of 10:90 and 90:10 are uniform down to ≤ 8°C, the lower limit of experiments. INSET: Grey dots are miscibility transition temperatures for ternary membranes of di(16:0)PC (DPPC), diPhyPC, and cholesterol, used with permission from (27). Interpolated colors range from ∼14°C at the far left to ∼47°C at the upper right. A larger version with a color scale is in Fig. S15; corresponding values are in Table S1 and S2. Grey dashed lines lie at ratios of 2:1 (triangle) and 3:1 (square) of cholesterol to di(16:0)PC. Points within ∼1 mol% of each line are plotted in the main panel.

### What structural elements lead to robust phase separation in membranes with sterol-lipids?

In isolation, Experiment 1 suggests a simple equivalence between sterol-lipids and individual molecules of sterols and lipids. Sterol-lipid A1 has only a single chain, implying that that the sterol-lipid might correspond to one sterol and one-half of a high-*T*_melt_ lipid, a ratio of ∼2:1 (or 67 mol% sterol). Indeed, miscibility transition temperatures for the binary membranes in Experiment 1 fall between a 2:1 and 3:1 ratio of sterol to high-*T*_melt_ lipid in corresponding ternary mixtures (Fig. 6). In this context, it makes sense that the binary membranes in Experiment 3 do not phase separate, even though they contain all three components necessary for liquid-liquid phase separation: chains with high-*T*_melt_ values, with low-*T*_melt_ values, and sterols. Both components of the membrane are sterol-lipids: Sterol-lipid A1 (PChemsPC) and Sterol-lipid D (OChemsPC) (Figs. 3 and S14). If both sterol-lipids roughly correspond to a high (2:1 to 3:1) ratio of sterol to lipid, then the membranes containing these molecules lie in the horizontal grey region in Fig. S15, far from any known miscibility phase transition in the corresponding ternary membranes.

However, any conclusion that sterol-lipids correspond to an exact ratio of sterol to lipid is not generalizable. Small structural details matter. Although Sterol-lipid A1 differs from Sterol-lipid A2 only in the connection between the sterol and the lipid (Fig. 2 and Fig. S16), Lo-Ld coexistence is robust in binary mixtures containing Sterol-lipid A1, and it is not robust in binary mixtures containing Sterol-lipid A2. Instead, in these mixtures, liquid domains appear in, at best, half of vesicles in slightly hypoosmotic conditions at 8°C, and no vesicles phase separate without an osmotic pressure difference (Experiment 4). The order of how the tails connect to the lipid backbone may also matter. A version of Sterol-lipid A1 with reversed tails produces an order parameter profile corresponding to a membrane containing only ∼20-25% sterol, possibly because the sterol does not orient with respect to the neighboring carbon chain in the same way that individual sterol and lipid chains orient (28). Nevertheless, sterol-lipids have order parameters characteristic of Lo phases (28), and they diminish the measurable heat capacity of the membrane’s solid-to-liquid transition, just as mixtures of sterols and lipids do. (29).

Turning our attention to the low-*T*_melt_ lipid in Experiment 1 (diPhyPC), what structural elements might allow it to phase separate so robustly from Sterol-lipid A (PChemsPC)? Empirically, coexisting liquid phases tend to persist to high temperatures in membranes containing diPhyPC (27, 30). The lipid diPhyPC is unusual in having multiply methylated chains, which give it an extremely low melting temperature, below-120°C (31). For comparison, another common low-*T*_melt_ lipid, di(18:1)PC (common name DOPC) melts at-18 ± 4°C (32). Indeed, when we replace diPhyPC with the alternative low-*T*_melt_ lipid, phase separation is less robust (Experiment 2). Again, a less robust transition means that, at best, liquid domains appear in half of vesicles in slightly hypoosmotic conditions at 8°C, and no vesicles phase separate without an osmotic pressure difference. This result implies the lipid mixtures in Experiment 2 lie close to a phase boundary, but not within it. Fig. S15 shows how this result is consistent with the outcome from Experiment 1. If Sterol-lipid A1 corresponds to a ratio of cholesterol to high-*T*_melt_ lipid between roughly 2:1 and 3:1 then a mixture of it with di(18:1)PC skirts the outer edge of the published 2-p-hase region for the corresponding ternary mixture (26).

### Alternative lipids and negative controls

The highly disordered chains of diPhyPC, which helped membranes phase separate in Experiment 1, render this lipid a poor choice when mixed with other sterol-lipids. No stable vesicles form in binary mixtures of diPhyPC with either Sterol-lipid B or C (BbGL-1 or CPG; Expts. 5 and 7, Fig. 3), likely because all three molecules are cone shaped. The problem is so severe that no vesicles formed from 100% Sterol-lipid B. diPhyPC’s conical shape is reflected in its large cross-sectional area (*e.g*., 80.5 ± 1.5 Å^2^ at 30°C, compared with 67.4 ± 1 Å^2^ for di(18:1)PC (33, 34); values for diPhyPC under different conditions are compiled elsewhere (33, 35)). A conical shape of diPhyPC is consistent with shallow membrane defects and low chain order in molecular dynamics simulations of diPhyPC membranes (36, 37).

We mitigated, but did not entirely solve, the problem of unstable membranes due to cone-shaped lipids by switching to the alternative low-*T*_melt_ lipid, di(18:1)PC. Transition temperatures of binary membranes of Sterol-lipid B and di(18:1)PC appear to lie below the lowest temperatures accessible in our experiments. Specifically, fewer than half of the vesicles phase separate in slightly hypoosmotic conditions at ∼8°C, and no vesicles phase separate without an osmotic pressure difference (Expt. 5, Fig. 3). Based on previously observed trends (38), we hypothesized that replacing di(18:1)PC with lipids with shorter chains would allow coexisting liquid phases to persist at higher temperatures. However, binary membranes of Sterol-lipid B and either di(16:1)PC or di(14:1)PC did not undergo robust liquid-liquid phase separation over our experimental temperatures. Similarly, binary membranes of Sterol-lipid C and the alternative low-*T*_melt_ lipid do not phase separate (CPG and di(18:1)PC, Expt. 8, Fig. 3). This was a surprise given that, at first glance, Sterol-lipid C looks most like a 1:1 ratio of a sterol and a two-tailed phospholipid. Moreover, membranes of 100% Sterol-lipid C exhibit high order (23). Chemical alterations to Sterol-lipid C that might allow it to phase separate in a binary membrane might include lengthening its chains to 16 carbons.

Last, we performed a battery of negative controls. At minimum, we expect a sterol-lipid to behave as a sterol, whether it has a saturated or unsaturated chain. When viewed by fluorescence microscopy, binary membranes comprised of cholesterol and phospholipids exhibit either one liquid phase (if the lipid has a low *T*_melt_) or solid-liquid coexistence (if the lipid has a high *T*_melt_) (39). Domains in a solid phase (which are often referred to as a gel phase) are typically non-circular, and they do not coalesce quickly when they collide. In line with these expectations from traditional binary membranes, we observe only one liquid phase in some of the binary membranes containing sterol lipids and coexisting solid and liquid phases in the others (Figs. 3 and 4). The last negative control in Fig. 4 (a mixture of Sterol-lipid D & di(16:0)PC) supports the assertion that preferential interactions between saturated lipid tails and cholesterol are necessary for liquid-liquid phase separation.

### Outlook

In addition to the sterol-lipids investigated here, others could be chemically synthesized. Structural components that could be varied include the length and unsaturation of the sterol-lipid’s acyl chain, the orientation of its sugar (α-linkage vs. β-linkage affects lipid orientations (40)), its charge (zwitterionic for Sterol-lipid A1, A2, and D, neutral for Sterol-lipid B, and charged for Sterol-lipid C), and the length of the linker between the sterol and the lipid backbone. Sterol lipids are also extracted from natural sources. For example, Huang et al. used an ACGal mixture of sterol-lipids (one of which is Sterol-lipid B) (22). One potential application of sterol lipids is in drug delivery (29).

Additional applications might lie in engineering minimal cell systems to function with only two classes of lipids (41); if one of those classes were a sterol-lipid, it would be interesting to test if the cell membrane phase separated. Micron-scale, reversible, liquid-liquid phase separation has previously been observed in more complex membranes of living cells (1, 42), where it has biological importance. For example, in yeast, phase separation of vacuole membranes promotes docking of lipid droplets during periods of nutrient restriction (43–45). Furthermore, yeast actively modify their lipid compositions to maintain their vacuole membranes near a miscibility transition (46). Our discovery that sterol-lipids can substitute for both a sterol and a high-*T*_melt_ lipid suggests a new role for sterol-lipids in biological membranes where they are found. In addition to functioning as a sterol (22, 23), sterol-lipids may enable liquid-liquid phase separation in membranes that otherwise lack sufficient saturated lipids. Our results also open new directions for researchers conducting research *in silico*. Our minimal membrane presents a new system for researchers to validate force fields and leverage recent advances in atomistic simulations of membranes undergoing liquid-liquid phase separation (47–49).

More fundamentally, 2-component bilayers that undergo liquid-liquid phase separation solve challenges in determining tie-lines in 3-component membranes. Tie-lines contain essential information: their endpoints reveal the ratio of lipids in each phase, and they provide a path along which chemical potentials of membrane components are equal and constant. In a system with only *two* components, tie-lines are straightforward to determine – only one tie-line is possible at each temperature. As a result, tie-lines in two-component systems are straightforward to traverse in the laboratory: the amount of only one type of lipid needs to be tuned. Two-component systems are also favored because their tie-lines are parallel at different temperatures. In contrast, in a lipid membrane with *three* components, multiple challenges arise:

I) the direction of each tie-line must be experimentally determined for every new set of molecules (39);
II)the directions of tie-lines found by measuring area fractions may only approximate the directions found by measuring mole fractions (50);
III) when the amount of only one component is changed, the system lies on an entirely new tie-line; and
IV) the direction of the tie-line changes with temperature (20).

As a result, when different labs seek to compare results in lipid bilayers, it is easy for them to verify that their systems lie along the same tie-line when they use 2-component membranes, and difficult when they use 3-component membranes. One caveat is that a membrane composed of only two kinds of molecules may not qualify as a 2-component membrane if the molecules are in different configurations that do not interconvert quickly on time scales of the experiment (4), and osmotic conditions must be reported.

## Conclusion

To summarize, we observe micron-scale, liquid-liquid phase separation in bilayer membranes of only two components, one of which is a sterol-lipid. The resulting phase diagram yields an upper miscibility critical point, a nearly textbook example showing two liquid phases at low temperature, critical behavior at the highest transition temperature, area fractions of phases consistent with the lever rule, and only one liquid phase otherwise. This clean system makes interpretation of results straightforward. Moreover, it gives us a window into what molecular features might be important. Independent of whether liquid-liquid phase separation occurs in a standard 3-component membrane or in a 2-component membrane with sterol-lipids, the membrane apparently must contain carbon chains with low order, chains with high order, and a sterol, where the sterols closely interact with the ordered chains. Although these attributes appear to be necessary, they are not sufficient. Structural details matter, such as the length of the linker between the sterol and the lipid backbone. We expect that these structural details will prove fertile ground for future all-atom simulations.

## Materials and Methods

### Chemicals

The lipid di(14:0)-cholesteryl-6-O-phosphatidyl-α-d-glucopyranoside (CPG) was synthesized as previously described (51). The lipids 1-palmitoyl(16:0)-2-cholesterylhemisuccinoyl-sn-glycero-3-phosphocholine (PChemsPC), 1-palmitoyl(16:0)-2-cholesterylcarbonoyl-sn-glycero-3-phosphocholine (PChcPC), 1-oleoyl(18:0)-2-cholesterylhemisuccinoyl-sn-glycero-3-phosphocholine (OChemsPC), cholesteryl 6-O-palmitoyl-β-D-galactopyranoside (BbGL-1), 1,2-dioleoyl-sn-glycero-3-phosphocholine (DOPC), dipalmitoyl(18:1)-phosphocholine (DPPC), diphytanoyl(16:0-4Me)-phosphocholine (diPhyPC), and dipalmitoyl(16:0)-phosphoethanolamine-N-(lissamine rhodamine B sulfonyl) (Rhod-DPPE) were purchased from Avanti Polar Lipids (Alabaster, AL). All lipids have only one length of acyl chain. To disambiguate the sterol-lipids in this study from others in the literature, ACGal, or 6-O-acyl-β-D-galactopyranoside, refers to a mixture of sterol-lipids with different acyl chains, one of which is BbGL-1 (22). Similarly, CAG, or cholesteryl-6-O-tetradecanoyl-α-D-glucopyranoside differs from BbGL-1 by having only 14 carbons (rather than 16) in its acyl chain and in the orientation of the sugar (α-linked rather than β-linked); both are found in *H. pylori* (19). Sucrose (ACS grade) was from Fisher Chemical, and vacuum grease (MOLYKOTE High Vacuum Grease) from DuPont. All chemicals were used as supplied, without further purification. Avanti Polar Lipids quotes purities at 99%.

### Electroformation

A total of 0.125 mg lipids was mixed in stock chloroform solutions. Lipid ratios included 0.8 mol% Rhod-DPPE to fluorescently label the liquid-disordered phase. The lipid mixture was spread at 60°C across an ITO-coated glass slide (Delta Technologies, Loveland, CO) with the side of a glass pipette, leaving a thin, even lipid film. The lipid-coated slide was then placed in a vacuum desiccator for ≥ 30 min to evaporate residual chloroform. Next, a chamber was formed as follows: vacuum grease was used to attach two 1-mm-thick Teflon spacers near the edges of the slide, and a second ITO-coated slide was placed on top of the spacers. The chamber was then filled with 100 mM sucrose in 18 MΩ-cm water, sealed with more vacuum grease, and placed in a 60°C oven. An AC field of 1 V at 10 Hz was applied across the two ITO surfaces at 60°C for 1 hr.

### Imaging

Vesicle solutions were imaged within 2 hrs. of production. They were removed from electroformation chambers, diluted, sandwiched between coverslips, sealed with vacuum grease, and imaged on a home-built temperature-controlled stage with a Nikon Eclipse ME600L upright epifluorescence microscope and a Hamamatsu C13440 digital camera.

Dilution entailed mixing 20 µL of vesicle solution with 130 µL of 100 mM sucrose (for equal osmotic pressures) or, more commonly, 80 mM sucrose (for a slight osmotic difference that minimized vesicle tubulation). Newly electroformed vesicles containing sterol-lipids are flaccid, unlike the ternary membranes they are designed to replace. Although membranes in these flaccid vesicles can phase separate, tubules and other non-spherical shapes often appear with changes in temperature, making transitions difficult to measure (Fig. S1). Tubulation may be exacerbated in our experiments because the constituent lipids likely have conical shapes and because the lipid mixtures do not include free sterols, which can mitigate membrane asymmetries via flip flop (52). To address this problem, we applied a small osmotic pressure difference sufficient to prevent most tubulation (100 mM sucrose inside, measured as 96 ± 3 mOsm on a Wescor VAPRO 5520 vapor pressure osmometer, and 80 mM outside, measured as 78 ± 3 mOsm). At higher concentration differences (100 mM inside, 50mM outside), some vesicles cyclically ruptured as in (53) and *T*_mix_ was ∼3°C higher. Membrane tension, whether due to osmotic pressure or other sources, is known to shift *T*_mix_ (54–56).

### Measuring *T*_mix_

To measure miscibility transition temperatures (*T*_mix_) of membranes, vesicles were cooled stepwise from 40°C to 8°C. Images of ∼100 vesicles were collected from multiple fields of view every 2°C near the transition temperature and at least every 4°C otherwise. At *T*_mix_, domains on each vesicle appear all at once (not through time). To confirm reversibility of the transition, temperature was incrementally raised back from 8°C to 40°C. The percentage of phase-separated vesicles at each temperature was manually recorded by a single user.

The most likely source of uncertainty in Fig. 6 is that electroformed vesicles have variations in size and excess area that become relevant when osmotic pressure is applied (Fig. S17). In addition, slight variations in lipid composition occur from vesicle to vesicle (57). In vesicles containing cholesterol, variation in asymmetry in lipid number between the two leaflets of the bilayer are mitigated by sterol flip-flop, whereas flip-flop rates of sterol-lipids should be lower because of their larger hydrophilic headgroups.

To find *T*_mix_, a non-linear least squares regression was used to fit a sigmoidal curve with the form: *Percent phase Separated* = *Max* (1 – (1 + exp (–(*T* – *T*_mix_) / w)))^-1^), where *Max* is the maximum percent of vesicles that phase separate, *T* is temperature, and *w* is the width of the sigmoidal curve. 95% confidence intervals were calculated from variances of fit parameters.

## Author Contributions

H.Q.N. and J.G.H. synthesized the CPG sterol-lipid. K.W. and S.L.K. designed the subsequent research plan. K.W. performed research and analyzed data. K.W. and S.L.K. wrote the manuscript and produced the figures.

## Acknowledgements

S.L.K. acknowledges funding from NSF MCB-1925731 and NSF MCB-2325819 and J.G.H. acknowledges NIH R01GM090262, NSF CHE-0196482, NSF CRIF Program (CHE-9808183), and NSF OSTI 97-24412. In addition, J.G.H. gratefully acknowledges support from the National Science Foundation for independent research and development during her tenure as Division Director of Chemistry.

## Supplementary Information for

### This file includes

Captions for Movie S1 and Movie S2

Figures S1 to S17

Table S1 to S2

#### Movie S1

Fluorescence micrographs of a giant unilamellar vesicle as temperature is decreased from 29.5°C to 28.5°C. At the highest temperature, the membrane is in one uniform phase. As temperature decreases, the membrane approaches a miscibility critical point characterized by small fluctuations in lipid composition. At temperatures below the critical point, liquid domains with fluctuating edges are observed. The vesicle is made from a mixture of 50:50 mol% PChemsPC and diPhyPC. Each image is a 200 ms exposure, and the time between frames is 5 seconds.

#### Movie S2

A set of fluorescence micrographs of a giant unilamellar vesicle as temperature is decreased from 40°C to 10°C. Below the miscibility transition temperature (frame 4), coexisting liquid phases are observed. When domains collide, they coalesce quickly. The vesicle is made from a mixture of 60:40 mol% PChemsPC and diPhyPC. Each image is a 200 ms exposure, and the time between frames is 30 seconds.

**Fig. S1:**
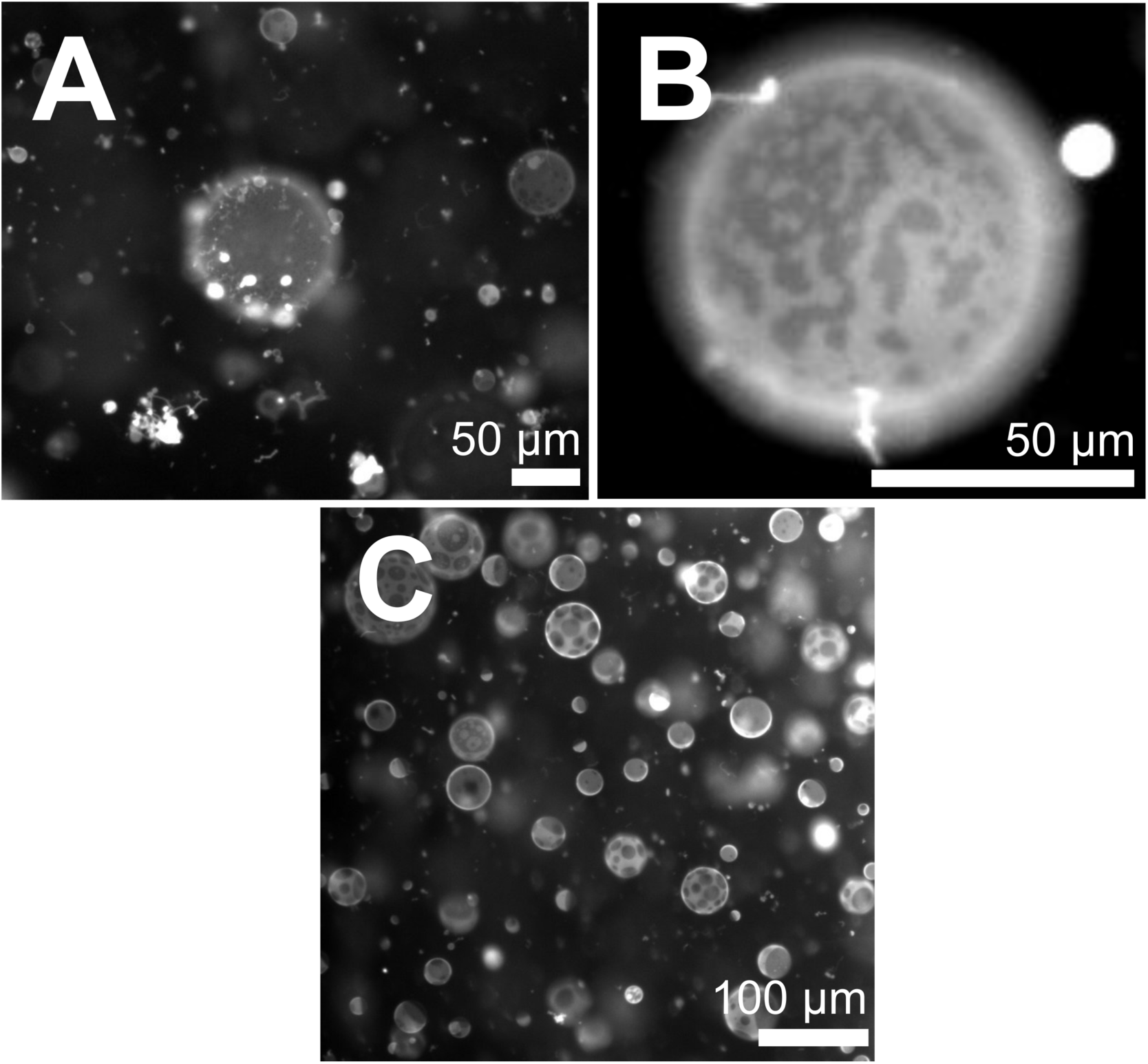
(A) Fluorescence micrograph of unilamellar vesicles composed of a mixture of 60:40 mol% PChemsPC and diPhyPC at 10°C. A 100 mM sucrose solution is inside and outside the vesicles. Tubules appear on surfaces of some membranes. (B) Shape fluctuations of membrane domains indicate miscibility critical behavior in a vesicle at 29°C made ofa mixture of 50:50 mol% PChemsPC and diPhyPC. This panel is from Supplementary Movie S1. (C) Wide, representative field of view of vesicles made from a mixture of 50:50 mol% PChemsPC and diPhyPC at 10°C. Vesicles were osmotically inflated: they were made in 100 mM sucrose and diluted in 80 mM sucrose.

**Fig. S2:**
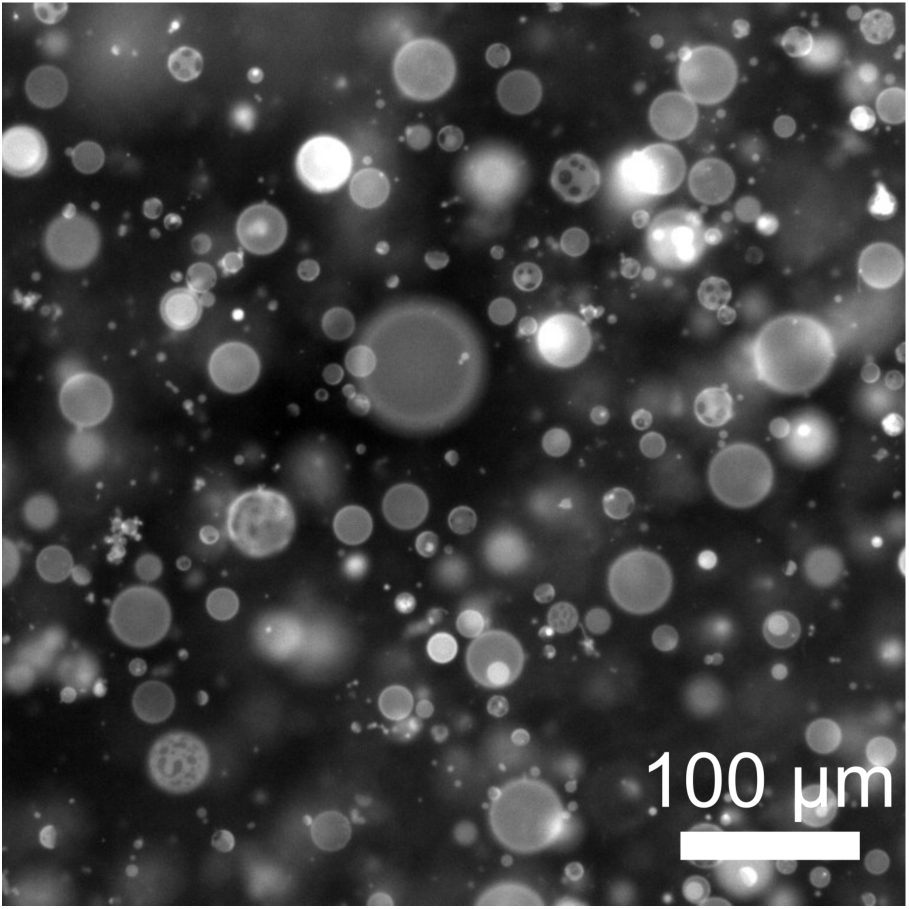
Wide, representative field of view of vesicles made from a mixture of 50:50 mol% PChemsPC and di(18:1)PC at 10°C. Some, but not all, vesicles display liquid-liquid phase separation. Vesicles were osmotically inflated: they were made in 100 mM sucrose and diluted into 80 mM sucrose.

**Fig. S3:**
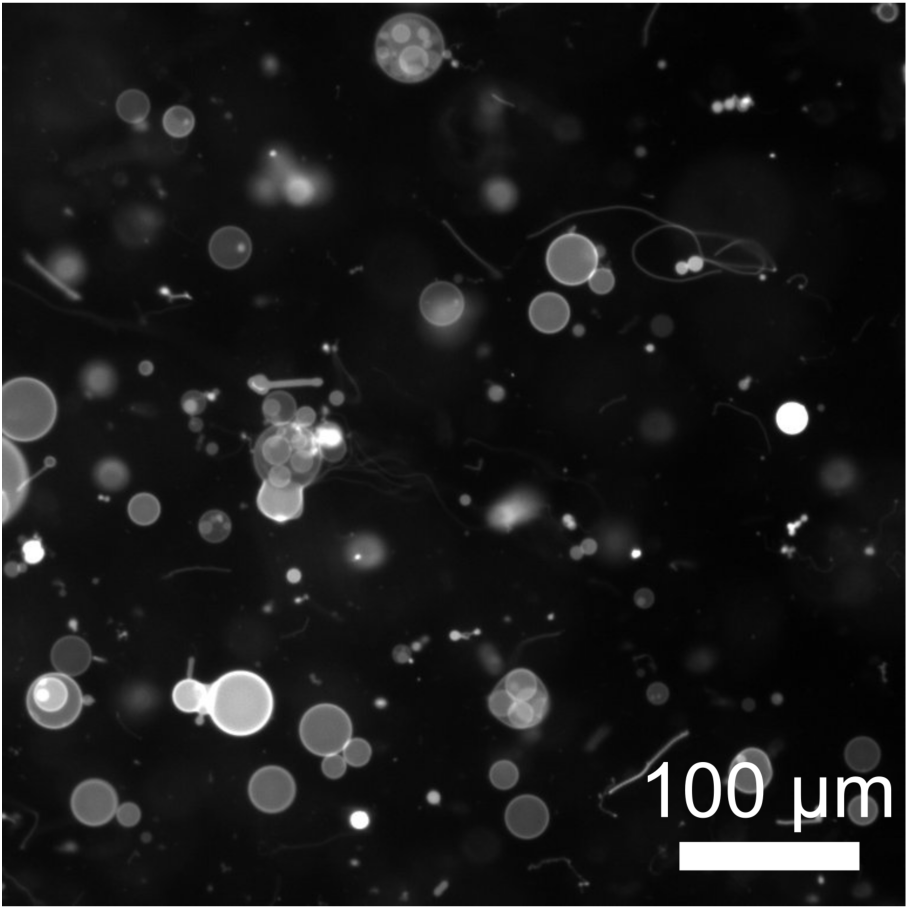
Wide, representative field of view of vesicles made from a mixture of 50:50 mol% PChemsPC and OChemsPC at 10°C. Additional structures are present including nested vesicles and tubules. Vesicles are osmotically inflated: they were made in 100 mM sucrose and diluted into 80 mM sucrose.

**Fig. S4:**
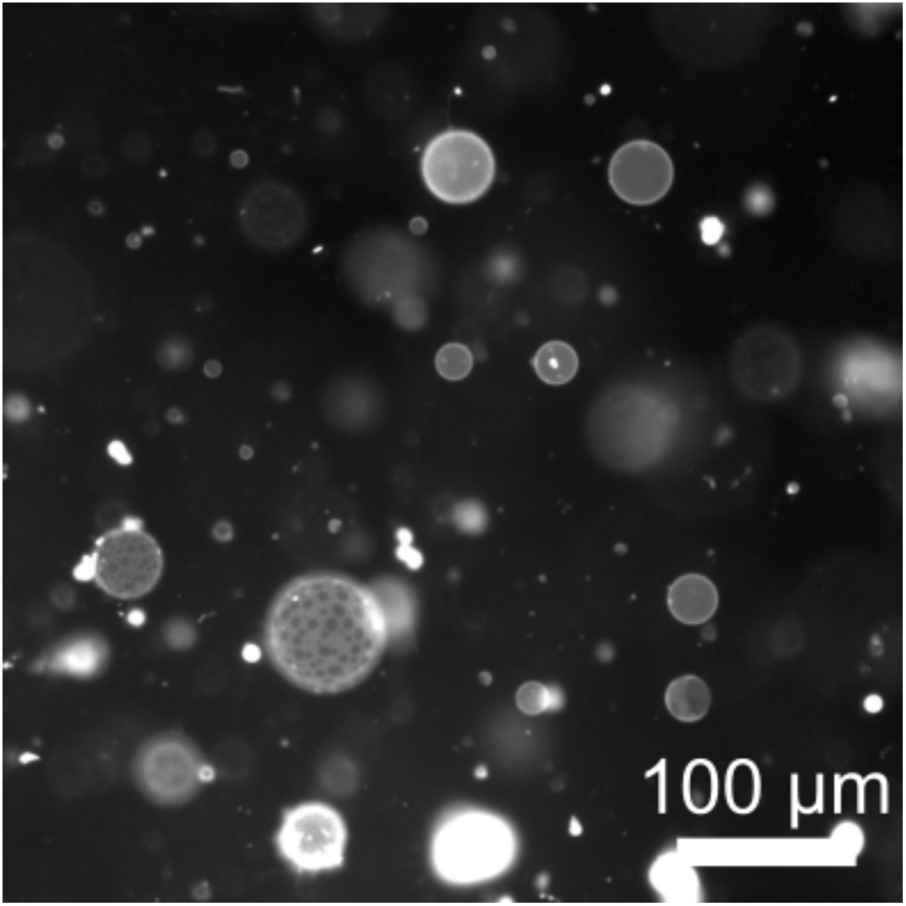
Wide, representative field of view of vesicles made from a mixture of 50:50 mol% PChcPC and diPhyPC at 10°C. Some, but not all, vesicles display liquid-liquid phase separation. Vesicles were osmotically inflated: they were made in 100 mM sucrose and diluted into 80 mM sucrose.

**Fig. S5:**
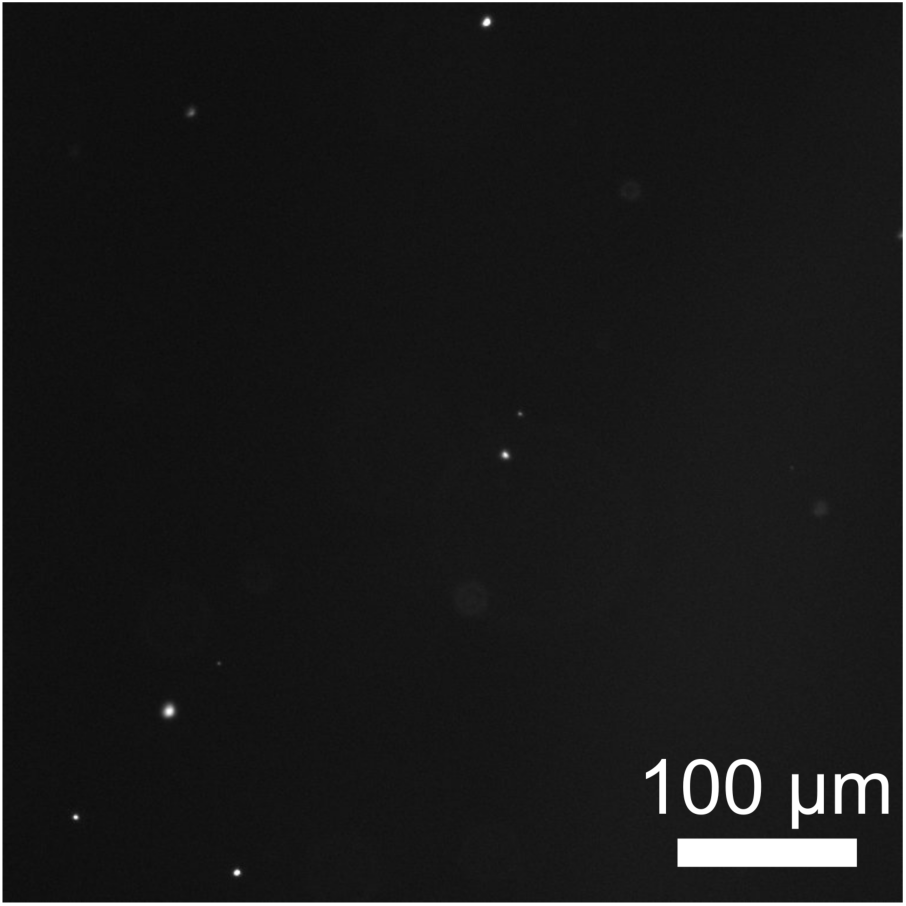
Wide, representative field of view showing that no vesicles were made from a mixture of 50:50 mol% BbGL-1 and diPhyPC.

**Fig. S6:**
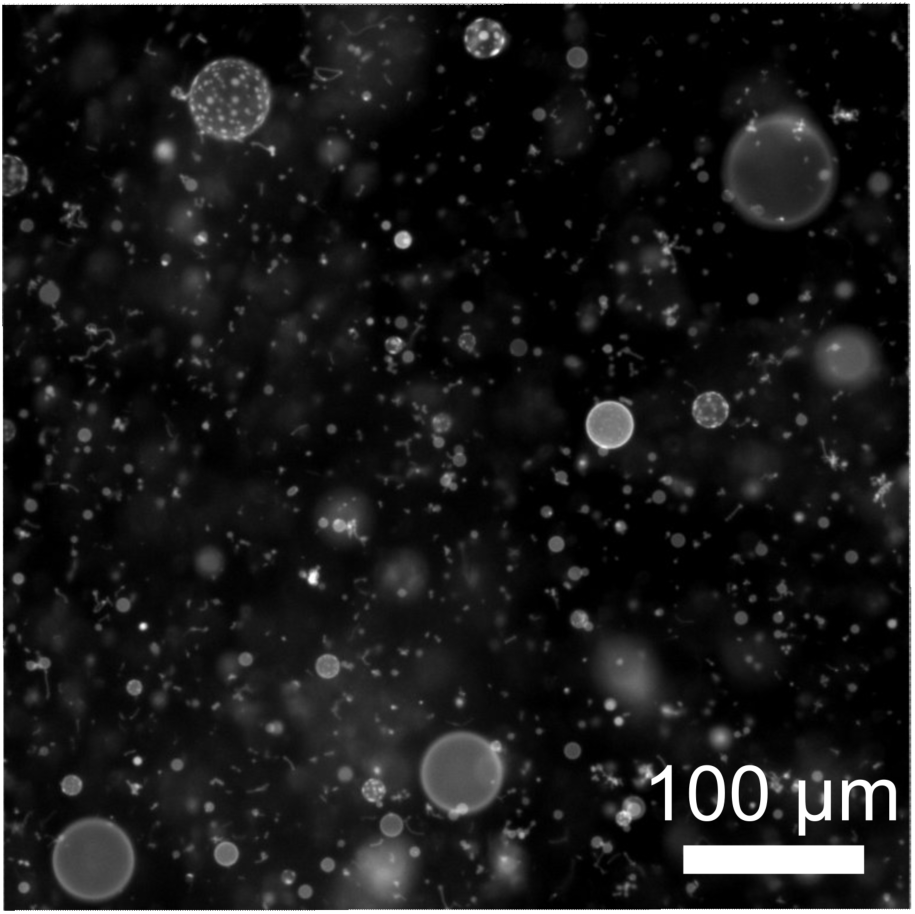
Wide, representative field of view of vesicles made from a mixture of 50:50 mol% BbGL-1 and di(18:1)PC at 10°C. Some, but not all, vesicles display liquid-liquid phase separation. Vesicles were osmotically inflated: they were made in 100 mM sucrose and diluted into 80 mM sucrose.

**Fig. S7:**
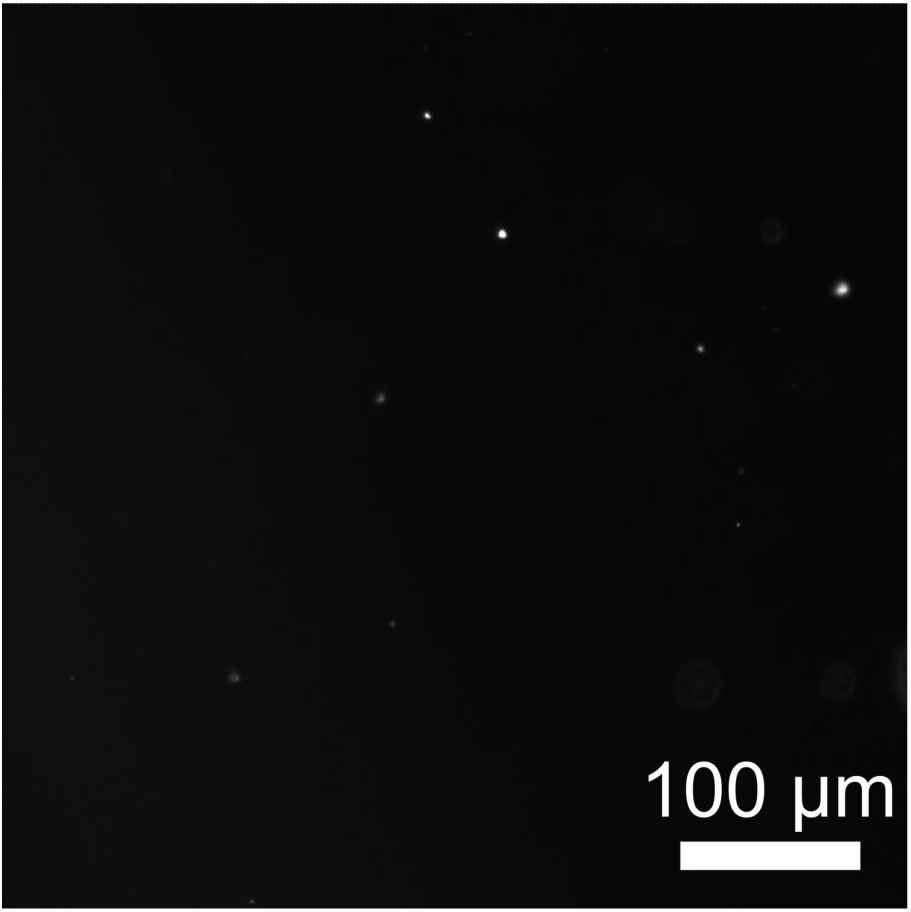
Wide, representative field of view showing that no vesicles were made from a mixture of 50:50 mol% CPG and diPhyPC.

**Fig. S8:**
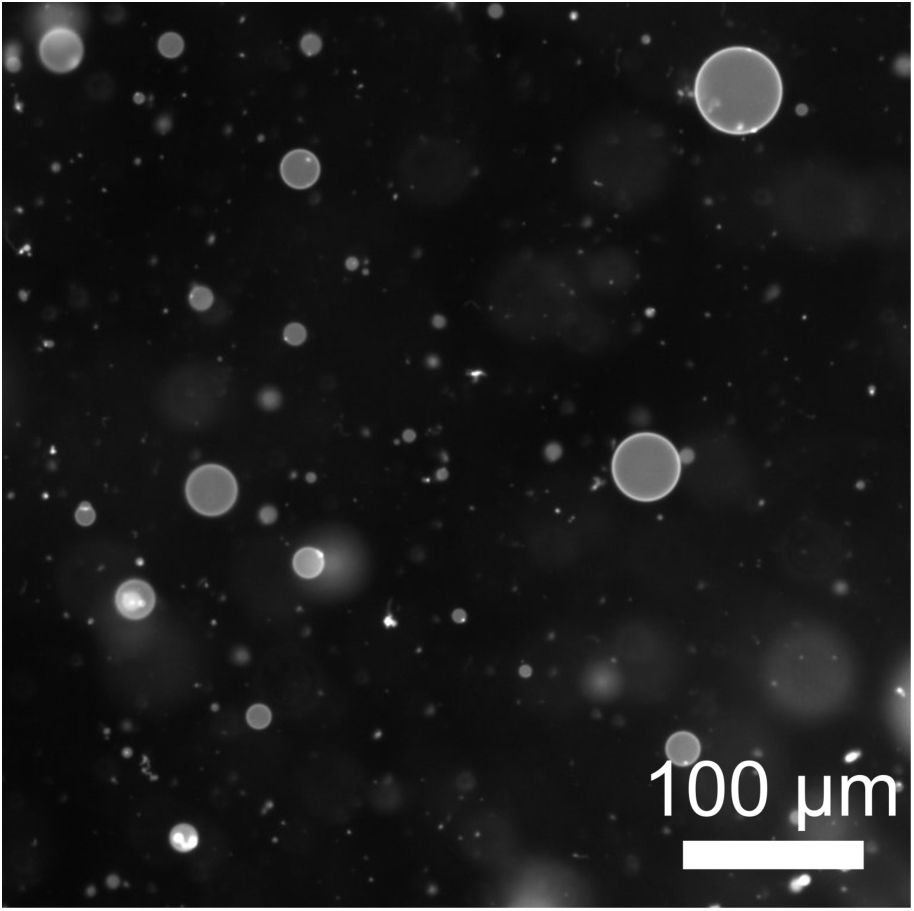
Wide, representative field of view of vesicles made from a mixture of 50:50 mol% CPG and di(18:1)PC at 10°C. Vesicles were osmotically inflated: they were electroformed in 100 mM sucrose and diluted into 80 mM sucrose.

**Fig. S9:**
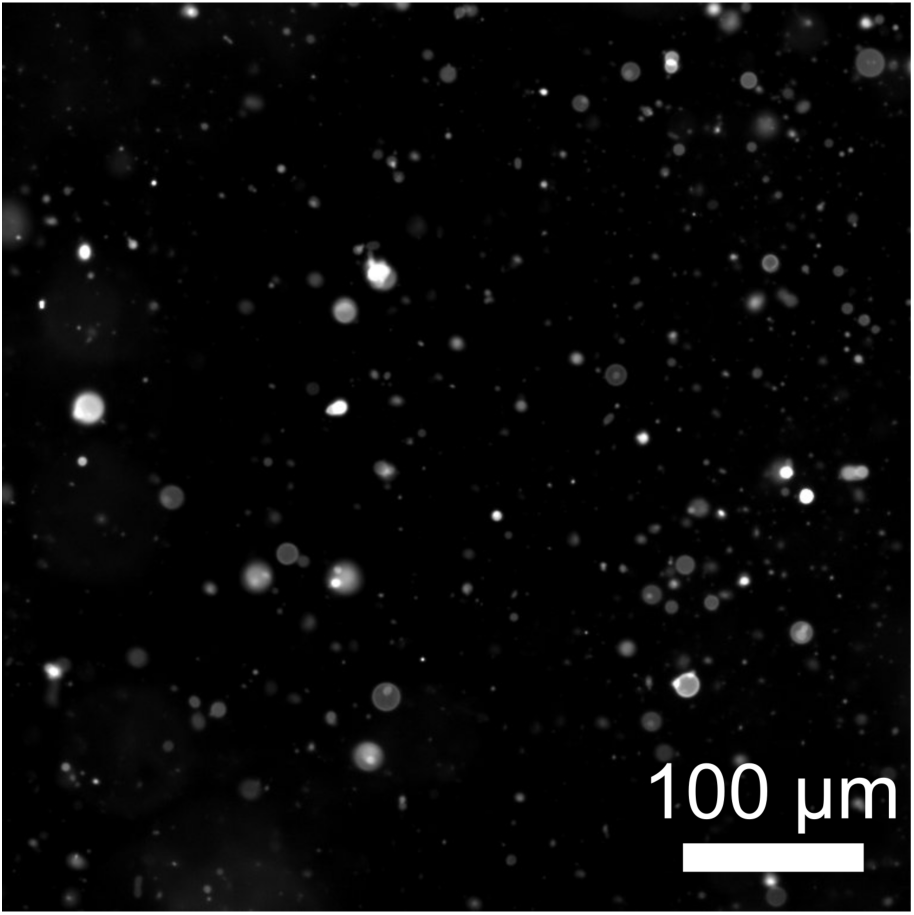
Wide, representative field of view of vesicles made from a mixture of PChemsPC and di(16:0)PC at 10°C. Vesicles were osmotically inflated: they were electroformed in 100 mM sucrose and diluted into 80 mM sucrose.

**Fig. S10:**
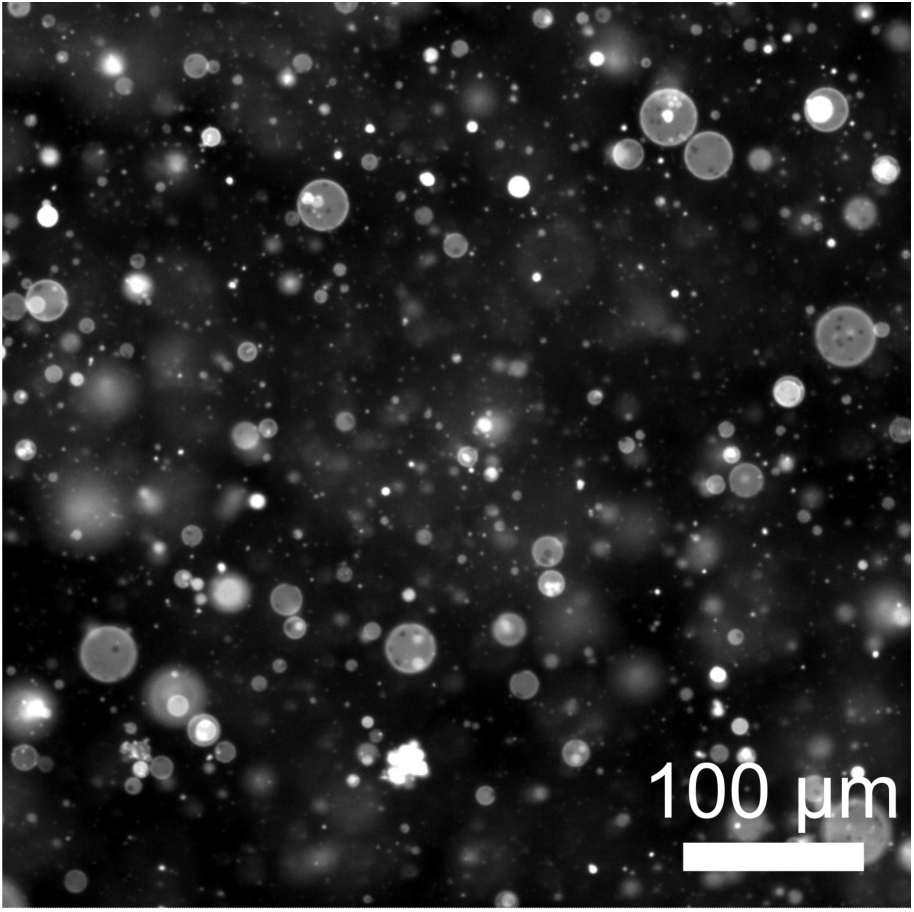
Wide, representative field of view of vesicles made from a mixture of OChemsPC and di(16:0)PC at 10°C. Most vesicles display solid-liquid phase coexistence. Vesicles were osmotically inflated: they were electroformed in 100 mM sucrose and diluted into 80 mM sucrose.

**Fig. S11:**
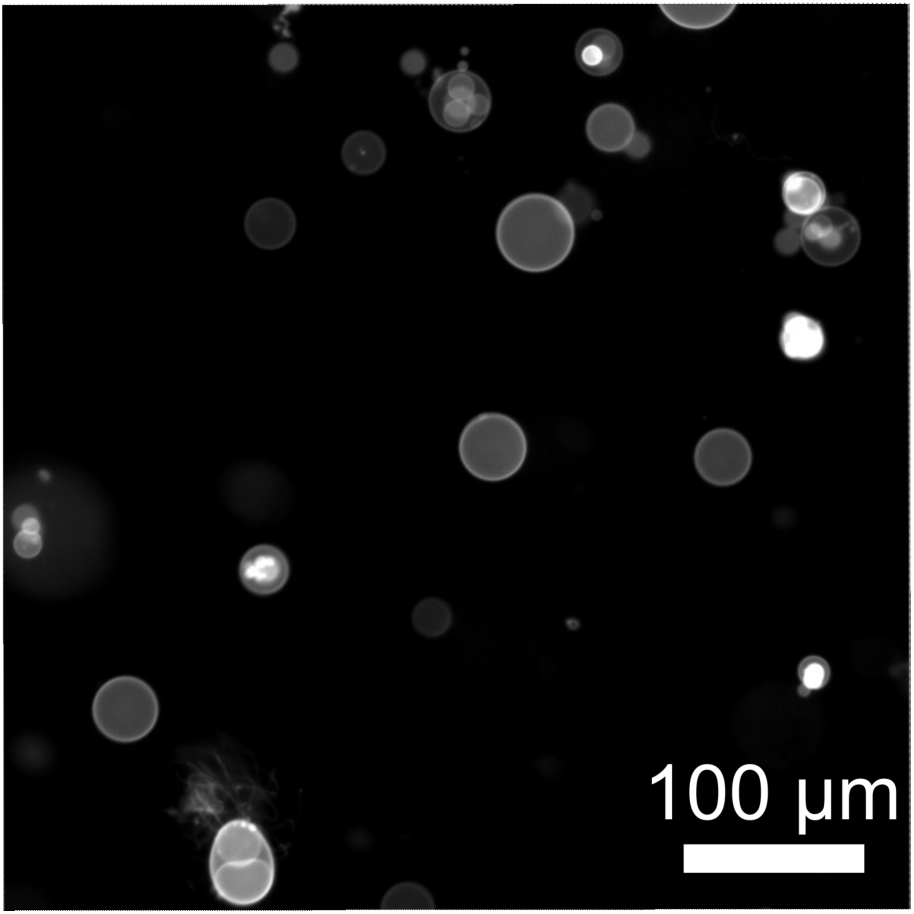
Wide, representative field of view of vesicles made from a mixture of OChemsPC and diPhyPC at 10°C. Vesicles were osmotically inflated: they were electroformed in 100 mM sucrose and diluted into 80 mM sucrose.

**Fig. S12:**
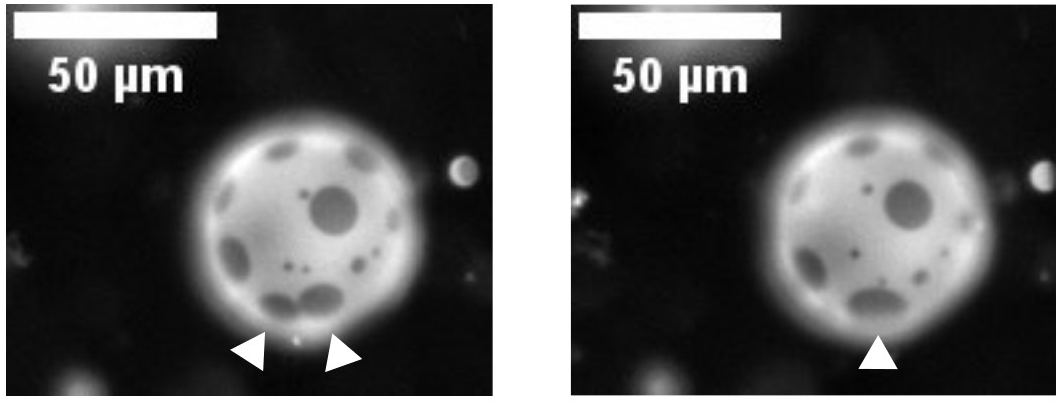
Liquid domains merging on the surface of a giant unilamellar vesicle made from a mixture of 60:40 mol% PChemsPC and diPhyPC. The images correspond to frame 11 (left) and frame 12 (right) of Movie S2. In both frames, the vesicle is at 10°C. The vesicles in this sample were electroformed in 100 mM sucrose and diluted into 80 mM sucrose. Each image is a 200 ms exposure, and the time between frames is 30 seconds.

**Fig. S13:**
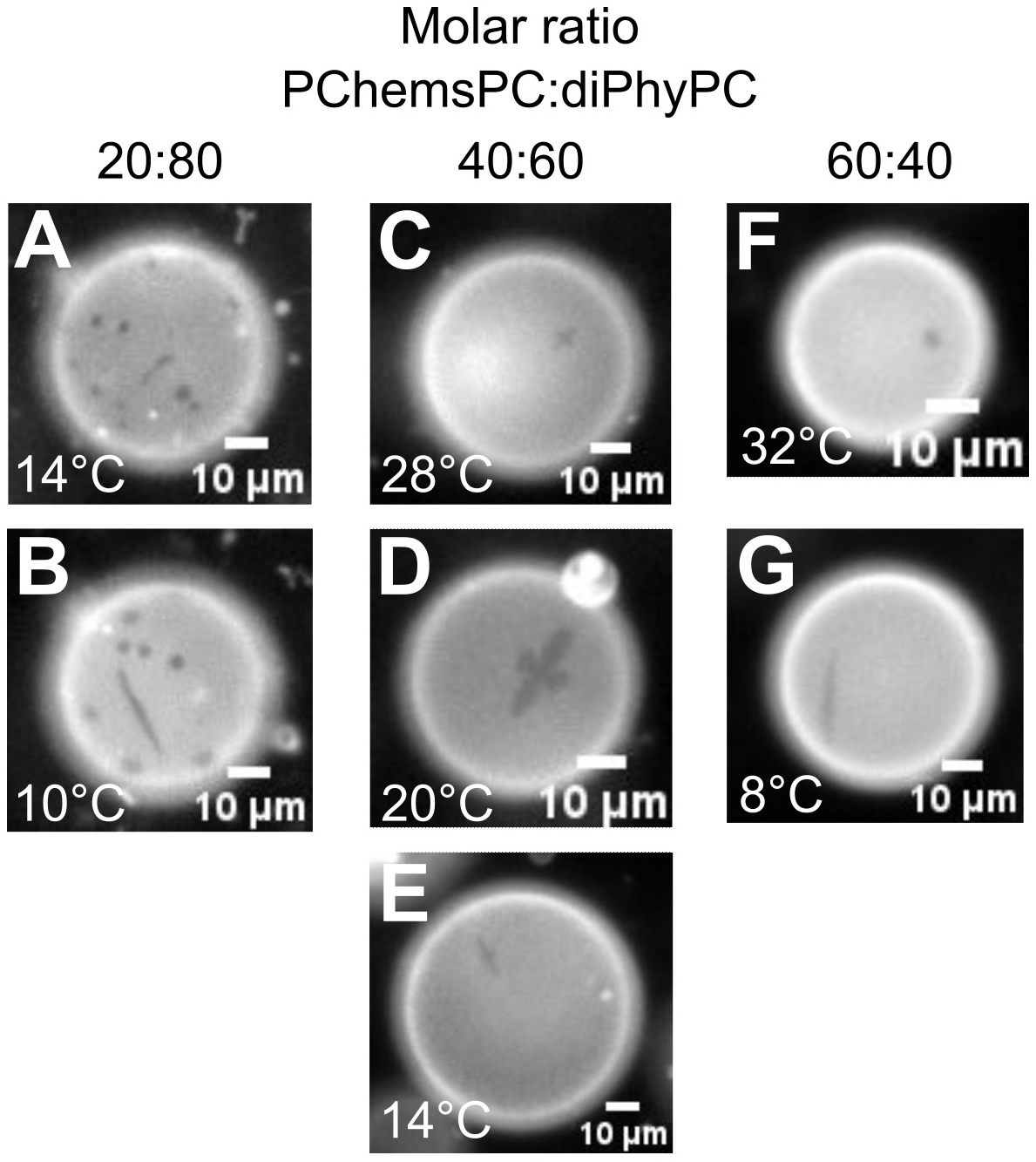
Examples of the largest “crystallites” observed in membranes of vesicles made from binary mixtures of PChemsPC and diPhyPC. In panels A and B, elongated crystallites in the center of the field of view coexist with circular, liquid domains. In Panels C to G, single crystallites are shown. Crystallites are over-represented in these images: on average, crystallites cover only ∼0.01% of the total area. This value represents an average of ∼1% of the area in an average of ∼1% of vesicles, with large sample-to-sample variation.

**Fig. S14:**
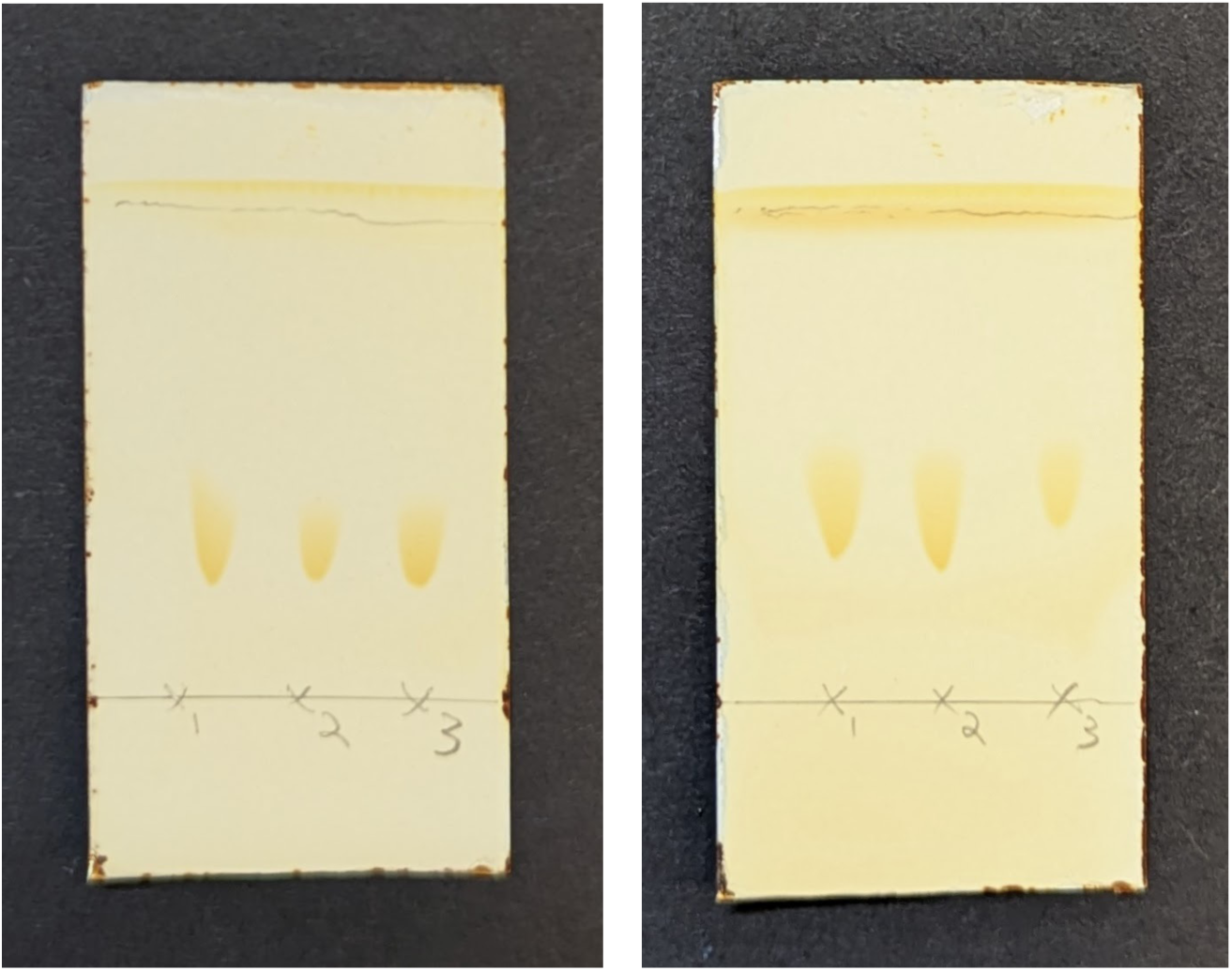
Triplicate thin layer chromatography spots for PChemsPC (left) and diPhyPC (right) as supplied by Avanti Polar Lipids. No impurities are detectable. The plates are 25 mm x 50 mm, and the solvent was 65:25:5 chloroform:methanol:water

**Fig. S15:**
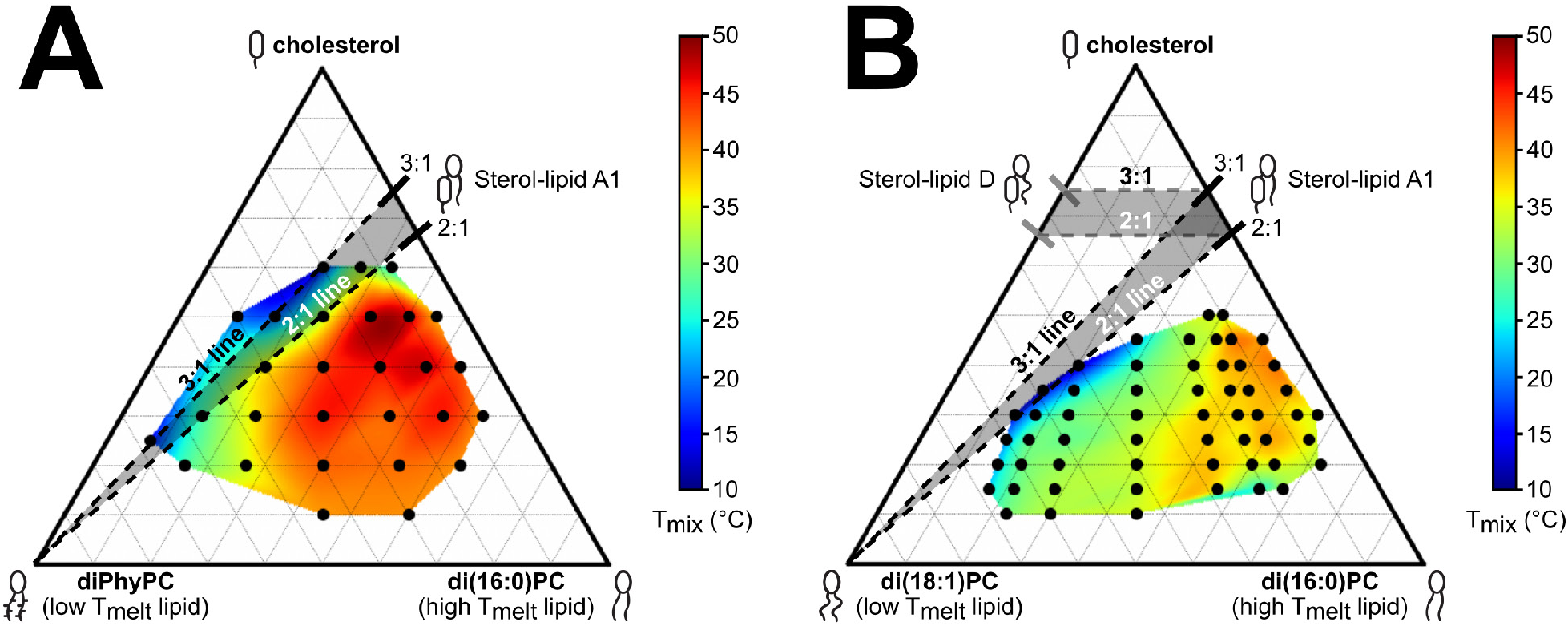
Miscibility transition temperatures at which coexisting liquid phases appear in ternary membranes of diPhyPC/di(16:0)PC/cholesterol measured down to 13°C (left panel) and di(18:1)PC/di(16:0)PC/cholesterol measured down to 10°C (right panel), used with permission from (27) and (26), respectively. The common name of di(16:0)PC is DPPC, and the common name of di(18:1)PC is DOPC. In the right panel, the angled shaded region corresponds to ternary membranes in which cholesterol and di(16:0)PC are maintained at a ratio between 2:1 and 3:1 and then mixed with pure di(18:1)PC. Similarly, the horizontal shaded region corresponds to ternary membranes in which cholesterol and di(16:0)PC are maintained at a ratio between 2:1 and 3:1 and then mixed with cholesterol and di(18:1)PC, which are also maintained at a ratio between 2:1 and 3:1.

**Fig. S16:**
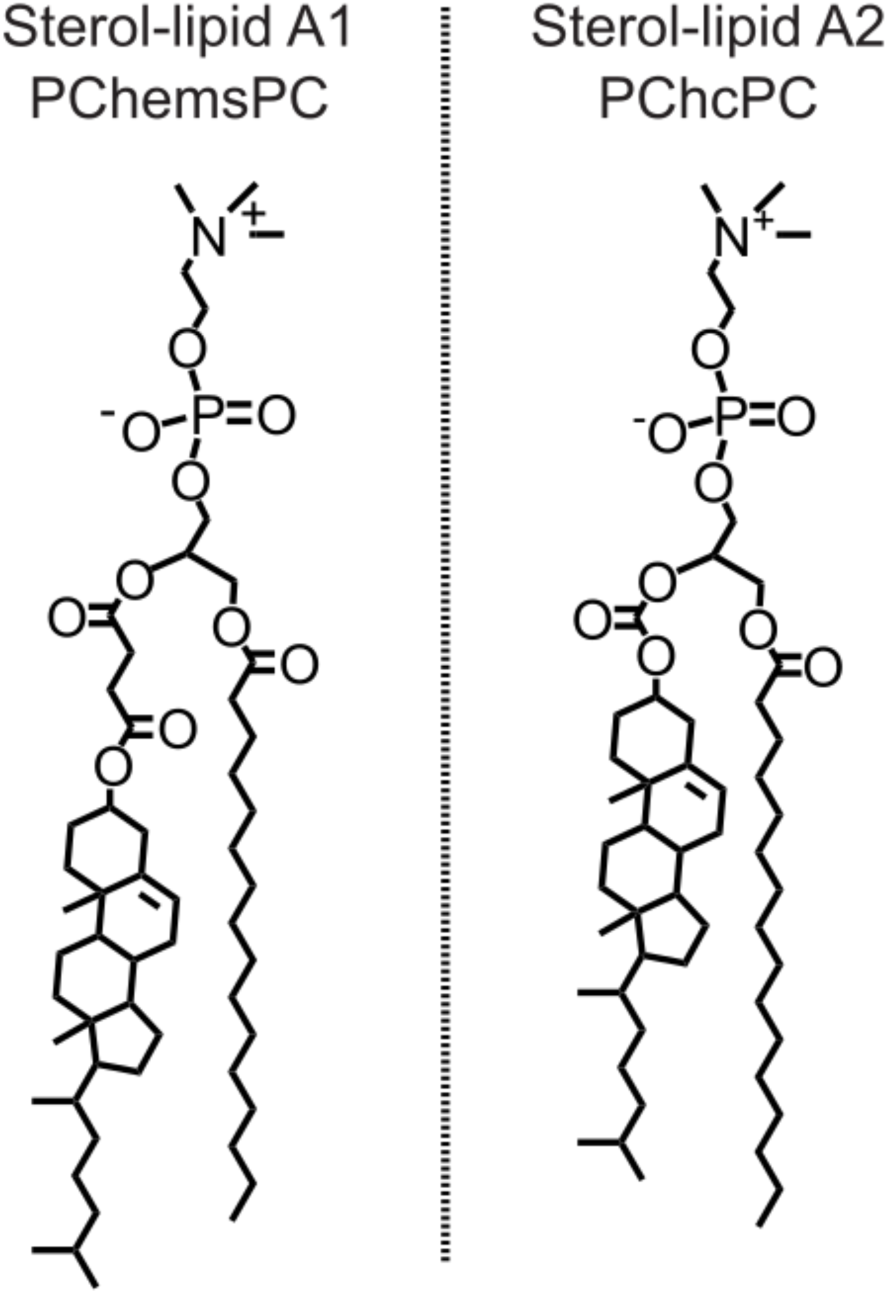
Comparison of the structures of Sterol-lipid A1 (PChemsPC, on the left) and Sterol-lipid A2 (PChcPC, on the right). The two sterol-lipids differ only in the connection between the sterol and the glycerol backbone.

**Fig. S17:**
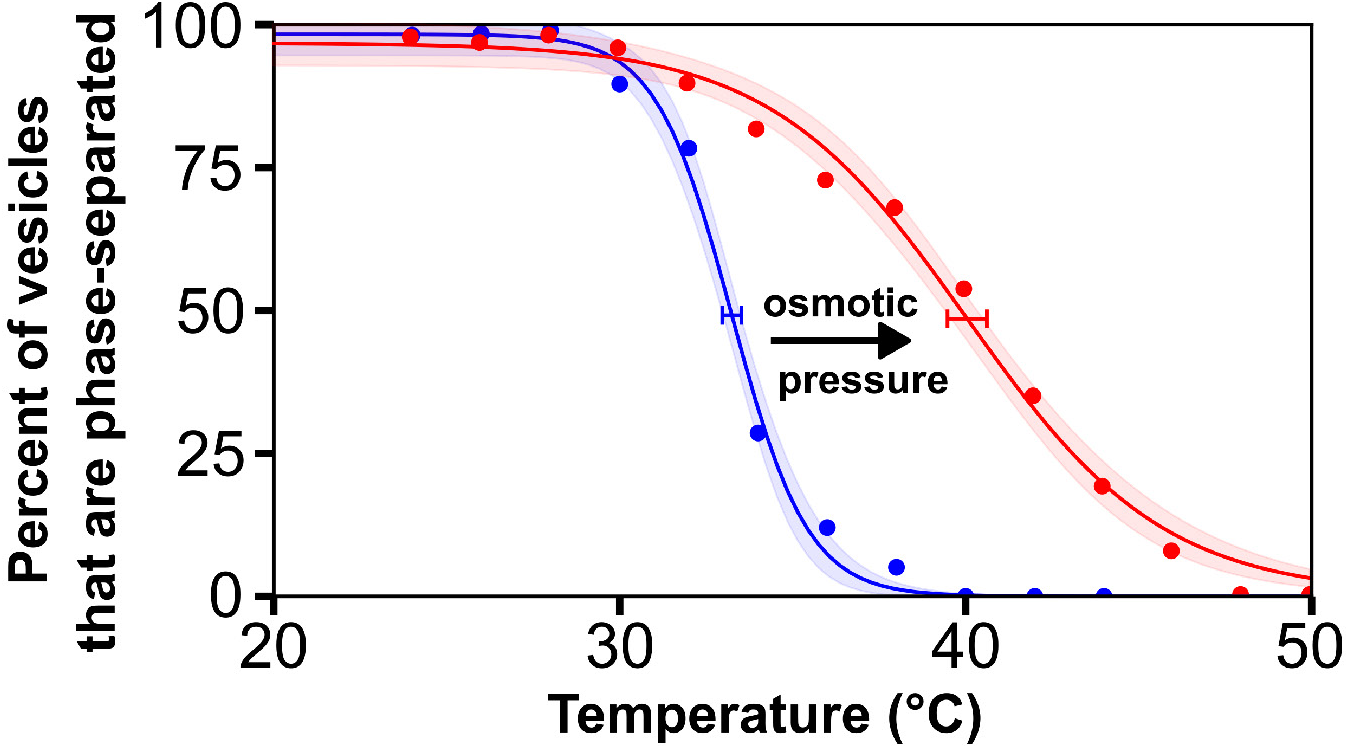
An osmotic pressure difference across a vesicle membrane increases and widens the miscibility transition temperature. The ratio of the lipids in the vesicle membranes is 20:40:40 mol% di(16:0)PC:diPhyPC:cholesterol; the common name if di(16:0)PC is DPPC. The full phase boundary for this ternary mixture is shown in Fig. S15. The ratio of lipids was chosen as a “worst case scenario” – it lies at a point in the phase diagram where the miscibility transition is very sensitive to changes in membrane composition (and, presumably, osmotic pressure differences). Vesicles were electroformed in 100 mM sucrose. The resulting vesicle solution was then split into two samples. The curve on the left corresponds to vesicles diluted in additional 100 mM sucrose. The curve on the right corresponds to vesicles diluted into 80 mM sucrose. Horizontal bars are drawn where 50% of vesicles have phase separated. The midpoint of each bar corresponds to the liquid-liquid phase transition temperature. The widths of the bars correspond to 95% confidence intervals of each fit. Other examples of increased membrane transition temperatures due to osmotic pressure differences, including in sugar solutions are found in (54–56). The increased width of the transition is likely due to variations in initial sucrose concentrations from vesicle to vesicle and in variations in vesicle sizes.

**Table S1:** Miscibility transition temperatures for vesicle membranes composed of Sterol-lipid A (PChemsPC) and diPhyPC. Uncertainties are 95% confidence intervals of the fit for individual experiments, as in Figure 5B. Vesicles were made in 100 mM sucrose and diluted into 80 mM sucrose. Miscibility transition temperatures could not be recorded for vesicles of 90:10 PChemsPC:diPhyPC because these membranes are uniform down to temperatures of ≤ 8°C, the lower limit of experiments. Preliminary experiments (right column) were conducted with initial manufacturing lots of PChemsPC and diPhyPC from Avanti Polar Lipids. Roughly one year later, independent, second manufacturing lots were ordered of both lipids, and three independent trials were conducted for each lipid ratio. Data from the three trials appear in Figure 6 of the main text.

| Molar ratio of<br>PChemsPC:<br>diPhyPC | Lipid Lots 2<br>Trial 1<br>Miscibility<br>transition | Lipid Lots 2<br>Trial 2<br>Miscibility<br>transition | Lipid Lots 2<br>Trial 3<br>Miscibility<br>transition | Lipid Lots 1<br>Miscibility<br>transition |
| --- | --- | --- | --- | --- |
| 10:90 | N/A | N/A | N/A | $7.9 \pm 0.1^{\circ}\text{C}$ |
| 20:80 | $11.5 \pm 0.6^{\circ}\text{C}$ | $13.1 \pm 1.6^{\circ}\text{C}$ | $13.5 \pm 0.1^{\circ}\text{C}$ | $20.0 \pm 0.6^{\circ}\text{C}$ |
| 30:70 | $21.3 \pm 0.4^{\circ}\text{C}$ | $21.4 \pm 0.4^{\circ}\text{C}$ | $14.3 \pm 0.7^{\circ}\text{C}$ | $28.8 \pm 0.9^{\circ}\text{C}$ |
| 40:60 | $28.6 \pm 0.4^{\circ}\text{C}$ | $21.7 \pm 0.5^{\circ}\text{C}$ | $25.3 \pm 0.4^{\circ}\text{C}$ | $30.8 \pm 0.9^{\circ}\text{C}$ |
| 50:50 | $24 \pm 0.6^{\circ}\text{C}$ | $21.4 \pm 0.3^{\circ}\text{C}$ | $21.7 \pm 0.4^{\circ}\text{C}$ | $28.8 \pm 0.9^{\circ}\text{C}$ |
| 60:40 | $18.3 \pm 0.2^{\circ}\text{C}$ | $23.3 \pm 0.4^{\circ}\text{C}$ | $24.2 \pm 0.3^{\circ}\text{C}$ | $25.0 \pm 1.5^{\circ}\text{C}$ |
| 70:30 | $16.1 \pm 0.2^{\circ}\text{C}$ | $21.2 \pm 0.3^{\circ}\text{C}$ | $22.8 \pm 0.3^{\circ}\text{C}$ | $21.7 \pm 2.0^{\circ}\text{C}$ |
| 80:20 | $15.8 \pm 1.5^{\circ}\text{C}$ | $18.8 \pm 1.5^{\circ}\text{C}$ | $17.2 \pm 0.7^{\circ}\text{C}$ | $19.8 \pm 1.2^{\circ}\text{C}$ |
| 90:10 | N/A | N/A | N/A | N/A |

**Table S2:**
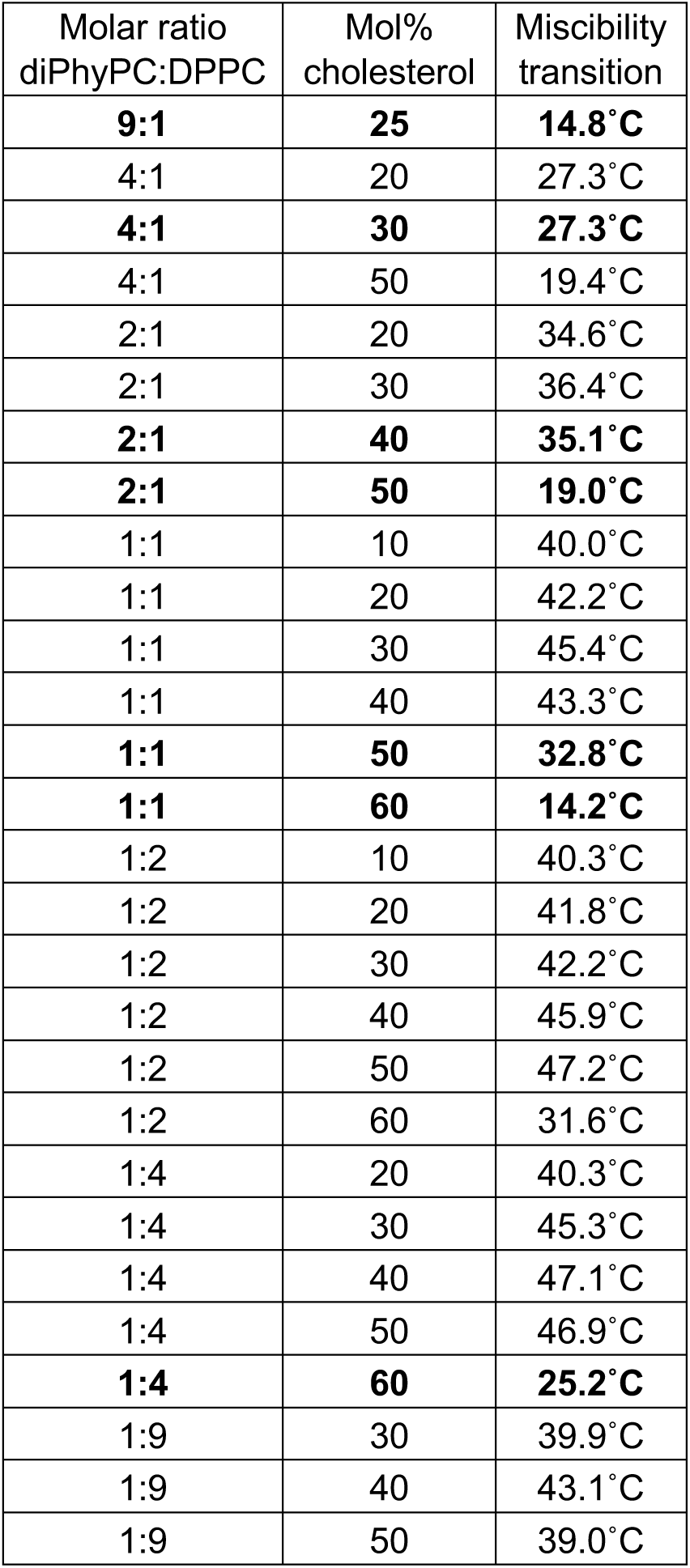
Miscibility transition temperatures for vesicle membranes composed of diPhyPC, DPPC, and cholesterol, used with permission from (27). Points in bold font are plotted in Figure 6.

